# Characterizing gene expression responses to biomechanical strain in an *in vitro* model of osteoarthritis

**DOI:** 10.1101/2021.02.22.432314

**Authors:** Anthony Hung, Genevieve Housman, Emilie A. Briscoe, Claudia Cuevas, Yoav Gilad

## Abstract

Osteoarthritis (**OA**) is a common chronic degenerative joint disease affecting articular cartilage and underlying bone. Both genetic and environmental factors appear to contribute to the development of this disease. Specifically, pathological levels of biomechanical stress on joints play a notable role in disease initiation and progression. Population-level gene expression studies of cartilage cells experiencing biomechanical stress may uncover gene-by-environment interactions relevant to OA and human joint health. To build a foundation for such studies, we applied differentiation protocols to develop an *in vitro* system of chondrogenic cell lines (**iPSC-chondrocytes**). We characterized gene regulatory responses of three human iPSC-chondrocyte lines to cyclic tensile strain treatment. We measured the contribution of biological and technical factors to gene expression variation in this system and, even in this small sample, found several genes that exhibit inter-individual expression differences in response to mechanical strain, including genes previously implicated in OA. Expanding this system to include iPSC-chondrocytes from a larger number of individuals will allow us to characterize and better understand gene-by-environment interactions related to OA and joint health.

## Introduction

Osteoarthritis (**OA**) is a chronic degenerative joint disease characterized by defects in articular cartilage integrity and alterations to underlying bone structure. OA is a major cause of disability in older adults and impacts approximately 300 million people worldwide^1^. There are currently no disease-modifying treatments for this painful disorder, and the specific pathogenic mechanisms of OA are still under investigation.

Genome-wide association studies (**GWAS**) have identified 86 genetic loci associated with OA risk^2^. Most of these loci fall within non-coding regions of the genome and have eluded functional characterization. Therefore, it remains unclear how associated genetic factors modulate OA onset and progression. One possibility is that regulatory changes in key structural and metabolic genes may modulate OA-related outcomes. Regulatory changes occurring in response to relevant environmental factors such as biomechanical stress may be particularly important. Indeed, gene expression studies have identified broad patterns of gene expression that differ markedly between healthy and osteoarthritic human cartilage^3,4^. These gene expression differences reflect activation of biological pathways associated with OA, suggesting that studies of gene regulation in cartilage, bone, and other skeletal tissues are valuable for understanding OA development and pathogenesis.

However, few studies have measured gene regulatory phenotypes in human skeletal tissues or cells. Even the Genotype-Tissue Expression (**GTEx**) Project, one of the largest efforts to examine gene expression variation across human tissues and cell types, does not include samples from either bone or cartilage^5^. This is partially due to the practical limitations and ethical issues associated with collecting healthy, high-quality cartilage and bone samples from human donors. Nevertheless, protocols to differentiate induced pluripotent stem cells (**iPSCs**) into OA-relevant cells, such as chondrocytes (the primary cells of cartilage), exist^6,7^, and these methods can circumvent some of the challenges associated with inaccessible primary tissues.

iPSC-derived cells also allow for the study of dynamic cellular responses to specific environmental conditions. It has become increasingly evident that studying gene regulation in disease-relevant states is crucial for understanding the genetic basis of disease^8^. Thus, numerous studies have begun identifying dynamic regulatory expression quantitative trait loci (**eQTLs**) in various cell types and contexts, including drug-induced cardiotoxicity^9^, cardiomyocyte differentiation^10^, vitamin D exposure^11^, and response to infection^12–16^. These studies highlight the merits of exploring gene regulation beyond steady-state conditions.

In OA, biomechanical stress is a particularly relevant environmental condition. Joint health deteriorates in response to excessive or insufficient amounts of mechanical loading^17–21^. Further, biomechanical factors may impact gene expression regulation in joint tissues and may interact with genetic factors to impact OA risk^22^. Such interactions are difficult to examine *in vivo*. However, iPSC-derived chondrogenic cells (**iPSC-chondrocytes**) provide an alternative system in which to study the effects of cyclic tensile strain (**CTS**), a type of controlled biomechanical stress regimen designed to induce OA-like phenotypes^23–26^. Thus, iPSC-chondrocytes offer an opportunity to study gene expression responses to OA-relevant states. Studies of iPSC-chondrocytes may also help uncover mechanisms through which OA-associated genetic loci modulate OA outcomes.

Both chondrocyte differentiation protocols and methods for inducing CTS *in vitro* existed prior to this study. Still, little is known about the suitability of this system for studies of gene regulatory dynamics. Therefore, we designed a study using human iPSC-chondrocytes to examine the combined effects of genetic variation and biomechanical stress on gene regulation during OA induction. Through this study, we evaluated whether the process of chondrocyte differentiation is robust to individual differences. We also ascertained whether iPSC-chondrocytes exhibit a robust gene expression response to CTS. Finally, we determined whether expanding the sample size of this experimental system might further improve our understanding of gene-by-environment interactions within OA and joint health.

## Results

We designed this study to determine whether iPSC-chondrocytes are a useful system for studying gene-by-environment regulatory interactions relevant to OA. First, we asked whether the efficiency of chondrocyte differentiation is similar in different individuals. Next, we evaluated the effects of cyclic tensile strain (CTS) on gene regulation in iPSC-chondrocytes to determine whether this system is suitable for studying gene regulatory effects on joint health. Finally, we estimated the contribution of sample and batch effects to variation in gene expression response to CTS, to assess the suitability of our iPSC-chondrocyte system for response eQTL mapping studies.

### Study design and data collection in the iPSC-based *in vitro* strain system

We used three human iPSC lines that were previously established and characterized as part of a panel of iPSCs derived from Yoruba individuals^27^. We differentiated the iPSCs along the chondrogenic lineage with an intermediate differentiation step into mesenchymal stem cells (**MSCs**; **Figure 1a)** using previously established protocols^6^. iPSC-derived MSCs exhibited phenotypes and cell surface marker expression patterns characteristic of primary MSCs^28^ (**Supplementary Figure S1**). Further, iPSC-chondrocytes showed a modest increase in collagenous extracellular matrix (**ECM**) production as compared to matched iPSC-derived MSCs (**Figure 1b; Supplementary Figure S2**).

**Figure 1:**
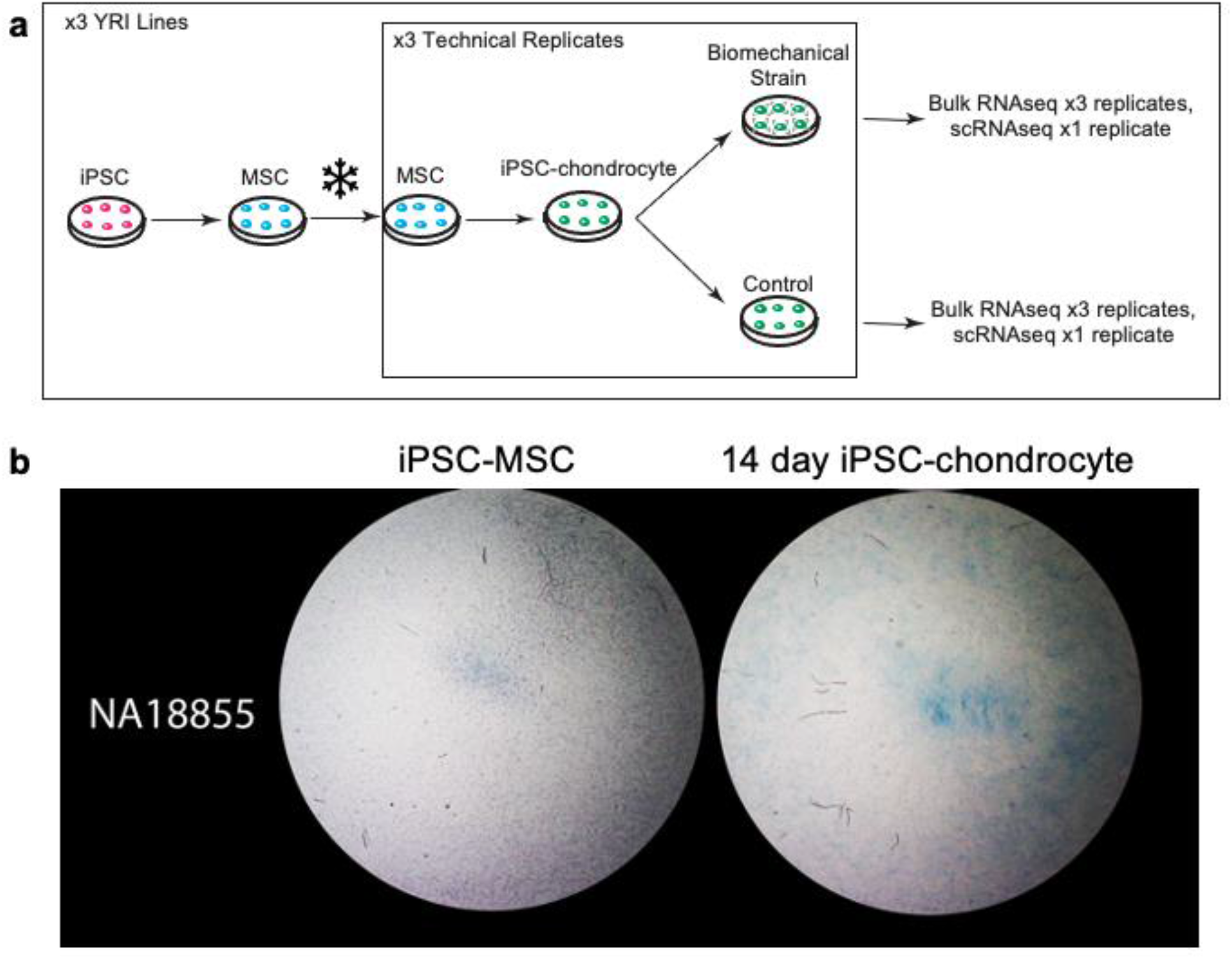
Description of *in vitro* biomechanical strain study design. **(a)** In our study design, iPSCs generated from three Yoruba individuals were first differentiated along the chondrogenic lineage with an intermediate differentiation step into mesenchymal stem cells (MSCs). iPSC-derived MSCs from each individual were cryopreserved. For each replicate of the experiment, iPSC-MSCs from the same cryopreservation batch were differentiated into iPSC-chondrocytes over a period of 14 days. Subsequently, we treated iPSC-chondrocytes from each individual with 24 hours of a CTS treatment that is known to induce an OA-like phenotype. We simultaneously kept a second, matched control set of iPSC-chondrocytes in the same incubator for the same period of time, but without CTS treatment. We performed three technical replicates of this experiment, starting with MSCs from the same cryopreservation batch. Following strain-treated and control conditions, we extracted bulk RNA from all biological and technical iPSC-chondrocyte replicates and collected scRNA-seq data from one technical replicate from each cell line using the 10X Genomics Chromium Single Cell Gene Expression platform. **(b)** Representative images of Alcian blue staining of 14 day iPSC-chondrocytes and matched iPSC-MSCs demonstrating increased proteoglycan production in iPSC-chondrocytes. Images are cropped to show the central seeded area of wells of BioFlex Type I Collagen coated 6-well Culture Plate (seeded area diameter 25mm). Additional images for other cell lines are available in **Figure S2**.

We treated iPSC-chondrocytes from each individual with 24 hours of CTS, which is known to induce an OA-like phenotype in cartilage^23–26^; **Methods**). We simultaneously kept a second, matched set of untreated iPSC-chondrocytes in the same incubator for the same period as control. We performed three technical replicates of this experiment, starting with MSCs from the same cryopreservation batch and carrying out an independent differentiation of the MSCs to iPSC-chondrocytes in each replicate. We extracted bulk RNA from all biological and technical iPSC-chondrocyte treated and untreated replicates (n=9). We also collected single cell RNA sequencing (**scRNA-seq**) data from one technical replicate (n=3) using the 10X Genomics platform.

### iPSC-chondrocytes likely represent an early stage of chondrogenic development

As a first step in our analysis, we confirmed that iPSCs successfully differentiated to chondrogenic cells. Using standard staining protocols (**Methods**), we demonstrated that our cells produce glycosaminoglycan ECM, a hallmark of chondrogenesis (**Figure 1b; Supplementary Figure S2**). We also used our scRNA-seq data to address two major questions: First, what is the approximate proportion of iPSCs that differentiated into chondrogenic cells in each individual? Second, what is the relative maturity of iPSC-chondrocytes?

We expected that chondrocyte differentiation might result in heterogeneous populations of cells at different stages along the chondrogenic lineage and that iPSC-chondrocyte purity might differ across individual cell lines. We used scRNA-seq data to assess potential differences in differentiation efficiency among the three individuals. Following standardization and normalization (**Methods**), unsupervised clustering of the single cell data revealed three clusters of cells in our untreated control samples (**Figure 2a**). The proportion of cell membership in each cluster is comparable across individuals (**Figure 2b**). Five percent of cells from individual NA18855 fall into cluster 2, along with 8% of cells from NA18856 and 7% of cells from NA19160. Based on gene expression patterns, cluster 2 consists of cells that are most like chondrocytes; for example, these cells show high expression of the canonical chondrogenic marker *COL11A1*^29,30^ **(Figure 2c**; **Methods**; additional genes shown in **Supplementary Figure S3**). We found no substantial difference in *COL11A1* expression between individuals. Thus, it appears, based on staining and gene expression patterns, that iPSC-chondrocytes are undergoing chondrogenesis, and importantly, cell type composition does not differ substantially among the three individuals in this study.

**Figure 2:**
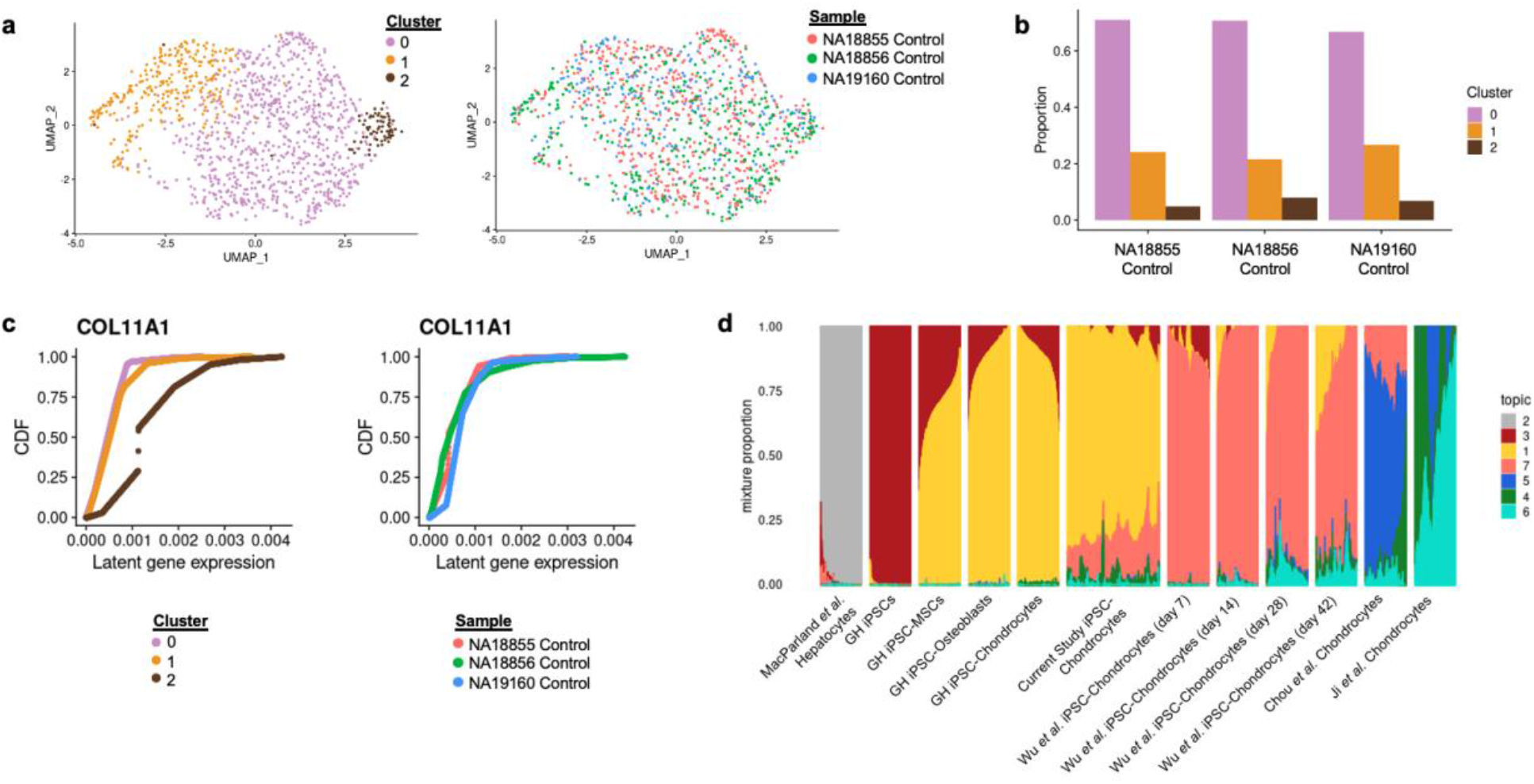
Characterization of cell type composition in iPSC-chondrocyte cultures. **(a)** Uniform Manifold Approximation and Projection (UMAP) of normalized and integrated single cell RNA sequencing data from control samples from all three individuals included in the study. (Left) UMAP colored by sample label. Cells from different samples are well-integrated. (Right) UMAP colored by Seurat cluster determined from normalized gene expression data. **(b)** Proportion of membership of cells from each individual in each Seurat cluster. The relative membership in each Seurat cluster is comparable between individuals. **(c)** Cumulative distribution function (CDF) of marginal distributions of latent gene expression of *COL11A1* determined through fitting a Poisson adaptive shrinkage model to raw gene expression counts in each sample. (Left) CDF curves colored by Seurat cluster. Cells in cluster 2 contain a higher latent gene expression of *COL11A1* on average. (Right) CDF curves colored by Individual. **(d)** STRUCTURE plot representing the relative proportional membership of single cells (columns) in 7 different topics in a topic model fit to scRNA-seq data derived from iPSC-chondrocytes collected in this study (“Current Study iPSC-Chondrocytes”); matched iPSCs, iPSC-MSCs, iPSC-osteoblasts, and iPSC-chondrocytes collected from a single individual (“GH iPSCs, GH iPSC-MSCs, GH iPSC-Osteoblasts, GH iPSC-Chondrocytes”); primary hepatocytes (“MacParland *et al*. Hepatocytes”); a time-course of iPSC-Chondrocyte pellet culture differentiation through the use of pellet culture (“Wu *et al*. iPSC-Chondrocytes (day 7 – day 42)”; and primary adult chondrocytes described in two separate publications (“Chou *et al*. Chondrocytes”, “Ji *et al*. Chondrocytes”). The ordering of individual cells within each study are determined through a one-dimensional tSNE algorithm applied to topic memberships of each cell. For each data source and cell type on the x-axis, a random subset of 800 cells is plotted with the exception of Current Study iPSC-Chondrocytes, for which all 1,815 cells are plotted.

In addition to discerning heterogeneity in our samples, we also evaluated the relative maturity of our iPSC-chondrocytes. To do this, we used topic modeling, analyzing our single cell data along with single cell datasets from multiple cell types, including primary adult chondrocytes (**Methods; Supplementary Note**). Topic modeling is an unsupervised classification approach that, when applied to single cell gene expression data, allows one to find recurring patterns of gene expression, or topics, present across a collection of cells. By allowing each cell to have grades of membership in multiple topics simultaneously, rather than assigning cells to only one cluster^31^, topic modeling can identify both discrete and continuous variation between cells.

A model fit with seven topics to the combined dataset shows that both iPSC-chondrocytes and primary chondrocytes are equally reliably distinct from unrelated cell types (e.g., hepatocytes). iPSC-chondrocytes retain a large proportion of gene expression patterns characteristic of iPSC-MSCs (topic 1), but they also possess certain gene expression patterns seen in adult primary chondrocytes and in iPSC-chondrocytes differentiated through a chondrogenic pellet (topics 4, 6, and 7) (**Figure 2d**). Topics 4-7 display relatively high levels of expression of several markers of chondrogenesis or cartilage fate, including *SOX9, SOX5, SOX6*^32^, and *COL9A1*^33^ (**Supplementary Figure S4**). Furthermore, differential expression analyses across topics identified type IX collagen gene *COL9A3*^33^ as highly occurring in topic 7 relative to all other topics (**Supplementary Figure S4**). Similarly, in topics 4 and 6, the chondrogenic marker *COMP*^34^ has a high occurrence relative to all other topics. Thus, we conclude that our iPSC-chondrocytes are likely in the early stages of chondrogenesis and are readily distinguishable from iPSCs and iPSC-MSCs.

### Analysis of bulk RNA sequencing data

After confirming that we can generate chondrogenic cells from iPSCs, we next sought to understand gene expression variation in this system. For this we focused on bulk RNA-seq data collected from all replicates. We generated an average of 22.3M raw reads per sample (s.d. 4M reads). We excluded one sample from further analyses because it displayed a particularly low percentage of mapped reads (**Supplementary Table S1**). We note that the mapped reads from this sample cluster as expected with other technical replicates from the same individual and treatment (**Supplementary Figure S5**), but we still excluded it because it failed standard QC metrics. We filtered the remaining data for lowly expressed genes and standardized gene counts with respect to library size (**Methods**).

As a first step of our analysis of the bulk data, we identified gene expression responses to strain treatment. We used the limma R package to fit a linear mixed model for each of the 10,486 expressed genes in the filtered bulk RNA-seq data, accounting for the random effect of experimental batch and the fixed effects for individual cell line, sex, treatment status, two factors of unwanted variation, and RIN score (**Methods**). Using this model, we tested for differential expression between treated and untreated cultures. At an FDR of 0.05, 987 genes are significantly differentially expressed (**DE**) between treated and untreated chondrogenic cells (**Figure 3a-b**; **Supplementary Table S2**).

**Figure 3:**
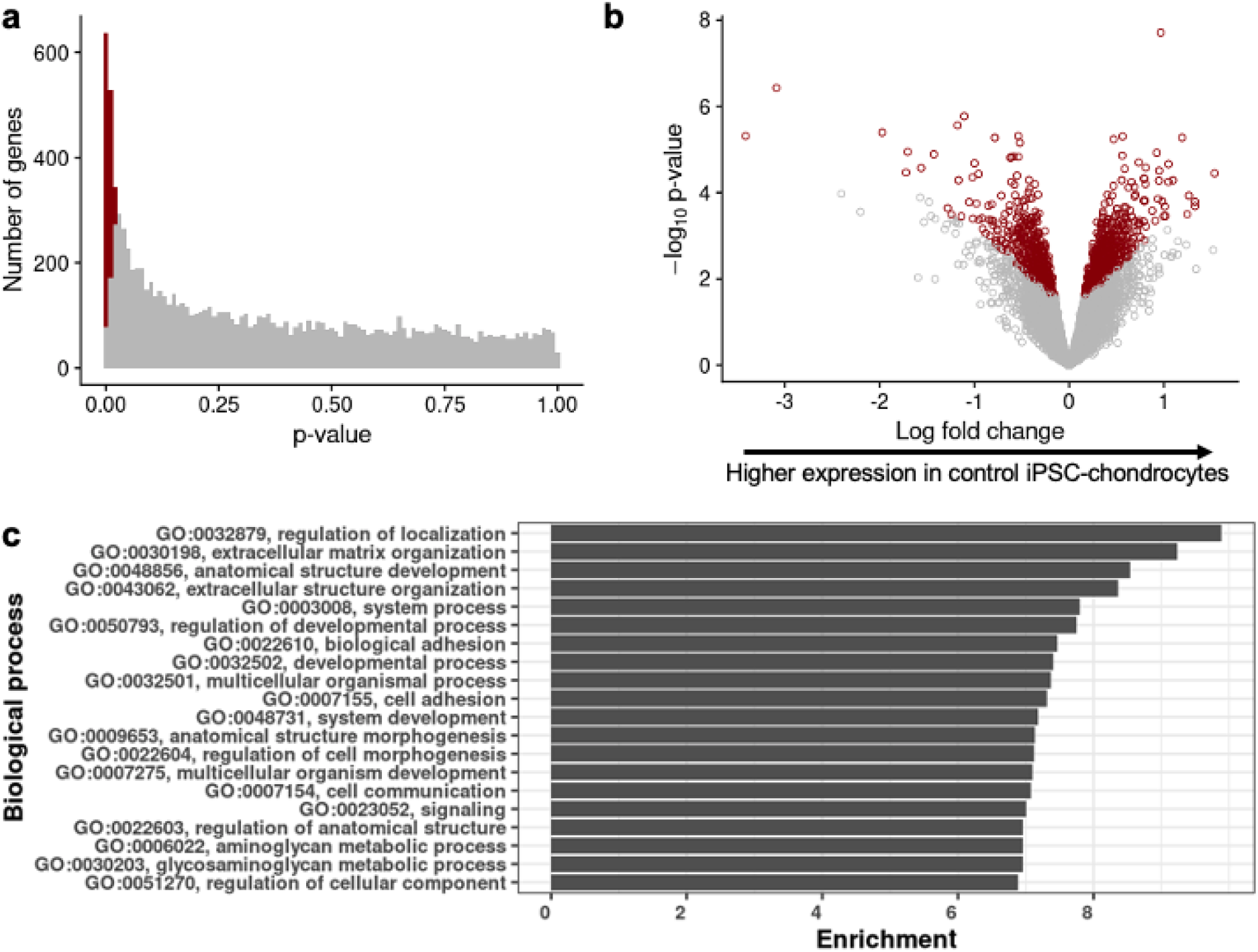
Results from linear mixed model differential expression (DE) analysis between treatment conditions. The linear mixed model used to conduct the analysis also accounted for the random effect of experimental batch and the fixed effects of individual cell line, sex, treatment status, two factors of unwanted variation, and RIN score. **(a)** Histogram of raw p values from DE analysis conducted independently for each gene between control and strain-treated conditions. Highlighted in red are genes with an FDR-adjusted p value < 0.05 (987 of 10,486 tested genes). **(b)** Volcano plot of –log10 raw p values vs log-fold change between treatment conditions. Highlighted in red are genes with an FDR-adjusted p value < 0.05. Genes plotted to the right of 0 on the x-axis represent genes with higher average expression in control iPSC-chondrocyte samples compared to strain-treated samples. **(c)** Top 20 Biological processes enriched among DE genes compared to background set of 987 genes. These GO biological process terms include those related to extracellular matrix organization and metabolism of extracellular matrix structure.

To evaluate the potential relevance of the DE genes to joint health and OA, we first considered their enrichment among gene ontology (**GO**) terms. The top 20 most highly enriched GO biological process terms include those related to ECM organization and metabolism of ECM structure (**Figure 3c**). These functions make intuitive sense given that ECM homeostasis is important for joint cartilage health; moreover, imbalances in this homeostasis are associated with OA^35,36^.

We also determined whether the DE genes may be overrepresented in gene sets previously implicated in OA. We examined results from the largest GWAS for OA susceptibility to date, which identified 64 independent significant associations with OA^2^. We used a Fisher’s exact test to assess enrichment of DE genes among a set of 553 genes located within 500 kb of the 64 associated loci. These 553 genes were also identified as having prior evidence of involvement in animal models of skeletal disease or human bone diseases^2^. We found that DE genes in our study are significantly enriched within the set of 553 genes previously associated with OA (p = 0.002; **Supplementary Table S3**).

Next, we evaluated results from a separate study, which profiled mRNA and protein samples in low-grade and high-grade osteoarthritic cartilage from 115 patients undergoing joint replacement^4^. Steinberg *et al*., 2019 found 409 genes with evidence of significant differential expression between patients with low-grade and high-grade osteoarthritic cartilage, at both the RNA and protein levels. Though causality is difficult to infer, this observation suggests that at least a subset of these genes is involved in OA cartilage degradation (**Supplementary Table S4**). A Fisher’s exact test reveals that our DE genes are also significantly enriched among this gene set (p = 0.02). Of note, the two genes that overlap between the two external gene sets, *LTBP3* and *LAMC1*, are also DE our study. *LTBP3* is a regulator of the TGF-ß signaling family, which plays roles in cartilage formation and development^37^. *LAMC1* has been identified as a blood-based biomarker for detecting mild knee OA, with lower RNA expression identifying mild OA^38^. Based on these GO and gene set enrichment results we concluded that the DE genes we identified between strain-treated and control iPSC-chondrocytes are relevant to joint health and OA.

Due to differential power, highly expressed genes are more likely to be detected as DE than lowly expressed genes in any RNA sequencing dataset^39^. Therefore, it is possible that the magnitude of expression of different genes in our data can explain our DE and enrichment results. To assess the robustness of our findings, we permuted the labels of treatment condition among our samples and re-performed DE and enrichment analyses a total of ten times (**Supplementary Table S5**). In nine permutations, we failed to identify ECM-related GO terms among the top 20 enriched terms (one permutation revealed two ECM-related GO terms). Further, we did not find any enrichment of DE genes within the two OA-relevant gene sets using permuted data. As our permuted data do not display the same enrichment patterns as our actual data, we concluded that our results are not due to differential power to detect DE.

### Certain gene expression responses to stress are heterogeneous between individuals

In our DE analysis, we focused on identifying inter-treatment differences in gene expression rather than inter-individual differences. Ultimately, we would like to use this system to study gene-by-environment interactions, which occur at the intersection of inter-treatment and inter-individual differences. A gene-by-environment interaction occurs when the magnitude or direction of gene expression response to an environmental stimulus is associated with an individual’s genotype at a particular locus. The sample size of this current study is far too small to detect gene-by-environment interactions. Still, we identified several genes that exhibit inter-individual differences in expression in response to CTS (**Figure 4**).

**Figure 4:**
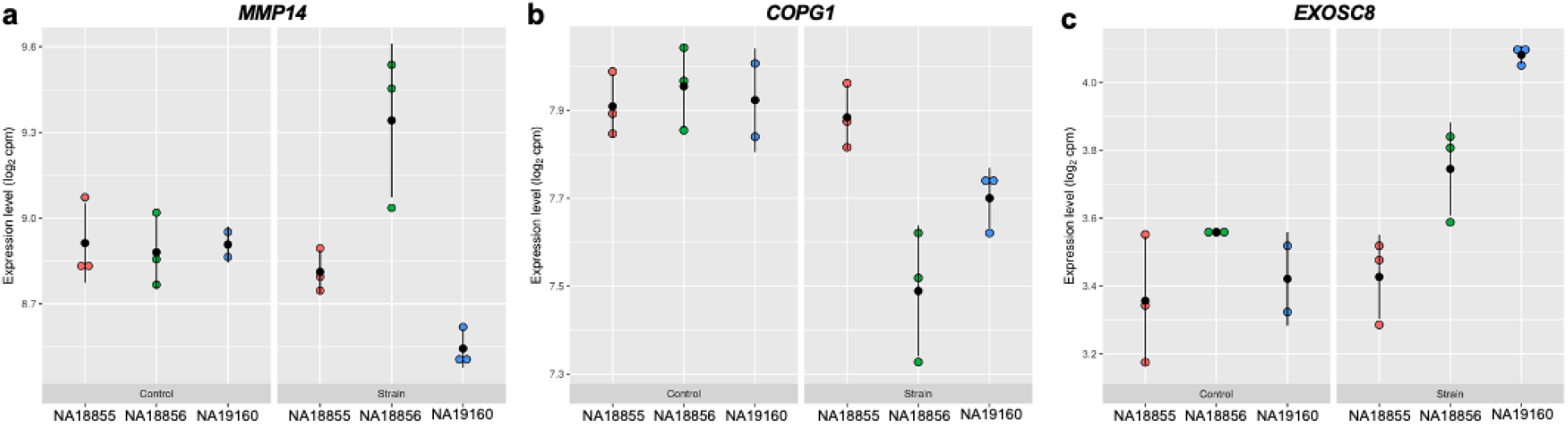
Examples of inter-individual differences in gene expression responses to cyclic tensile strain (CTS). **(a-c)** *MMP14, COPG1*, and *EXOSC8* each demonstrate inter-individual differences in gene expression in our dataset. Dot plots are the expression level (log2 cpm) of each gene in each sample, with each individual and treatment condition plotted separately. Lines represent +/-one standard deviation. For each candidate gene, expression is relatively consistent between individuals in the control condition but differs in magnitude or direction between individuals in the strain condition (in response to CTS). These heterogenous responses do not all involve differing magnitudes of changes in the same direction. In the case of *MMP14*, while NA18855 does not seem to respond to CTS through altering *MMP14* expression, NA18856 responds through upregulation of this gene and NA19160 responds through downregulation. If heterogenous responses to CTS such as these exist more broadly between individuals and are associated with genotypic differences, they should be identifiable in population-level eQTL studies using this experimental system.

For example, *MMP14* displays a different pattern of expression in each cell line before and after CTS (**Figure 4a**): *MMP14* expression remains constant between control and strain-treated NA18855 cells, is upregulated in strain-treated NA18856 cells, and is downregulated in strain-treated NA19160 cells. *MMP14* is expressed constitutively in adult joint cartilage and upregulated in diseased states^40^. In addition, the genes *EXOSC8* and *COPG1* (**Figure 4b-c**) are both involved in the formation of secretory vesicles originating from the Golgi complex. These genes also display differences in direction or magnitude of gene expression response to CTS between individuals. If heterogenous responses to biomechanical stress exist more broadly and are associated with genotypic differences, this experimental system will be able to identify them in population-level eQTL studies.

### Sources of variation in bulk RNA sequencing data

Thus far, our results show that CTS elicits a robust, OA-relevant gene expression response in iPSC-chondrocytes, and that, anecdotally, this response can differ between individuals. Next, we sought to more generally evaluate the utility of this system for studying the effects of genetic variation and biomechanical stress on gene regulation. Specifically, we considered dynamic expression quantitative trait loci (**dynamic eQTLs**), which are genetic variants associated with a change in gene expression in response to a treatment. For our system to be useful in mapping and identifying dynamic eQTLs, individual differences should drive a substantial amount of the variation in gene expression response to the treatment. To quantify the contribution of individual differences to gene expression variation in iPSC-chondrocytes, we estimated how much of this variation is attributable to technical and biological factors. Our study design allows us to do so, as we collected bulk RNA-seq data from three independent technical replicates of each cell line.

First, we evaluated the contributions of experimental variables to major axes of variation in our bulk RNA-seq data by performing a principal components analysis (**PCA**; **Supplementary Figures S6, S7; Supplementary Table S6**). Our results indicate that the primary source of gene expression variation is individual (regression of PC1 by individual, p = 1.34 × 10^−7^, regression of PC2 by individual, p = 4.61 × 10^−4^). The second largest source of variation in the data is sex (regression of PC2 by sex, p = 2.67 × 10^−4^), which is unsurprising given that our study included one female and two male cell lines. Although treatment shows a minor correlation with PC2 (R^2^ = 0.14), PC3 (R^2^ = 0.14), and PC4 (R^2^ = 0.3), none of these correlations are statistically significant.

Encouragingly, we did not find technical replicate (or ‘batch’) to be significantly associated with any of the first five PCs in the data. Nevertheless, we took advantage of our replicated experimental design to account for two factors of unwanted technical variation in the data^41^; **Methods**). Following this we observed that the top three sources of gene expression variation are individual (regression of PC1 by individual, p = 7.34 × 10^−8^, regression of PC2 by individual, p = 3.98 × 10^−4^), sex (regression of PC2 by sex, p = 1.79× 10^−4^), and treatment effect (regression of PC3 by treatment, p = 3.92 × 10^−2^), all three of which are significant (**Figure 5a-b, Supplementary Figure S8**).

**Figure 5:**
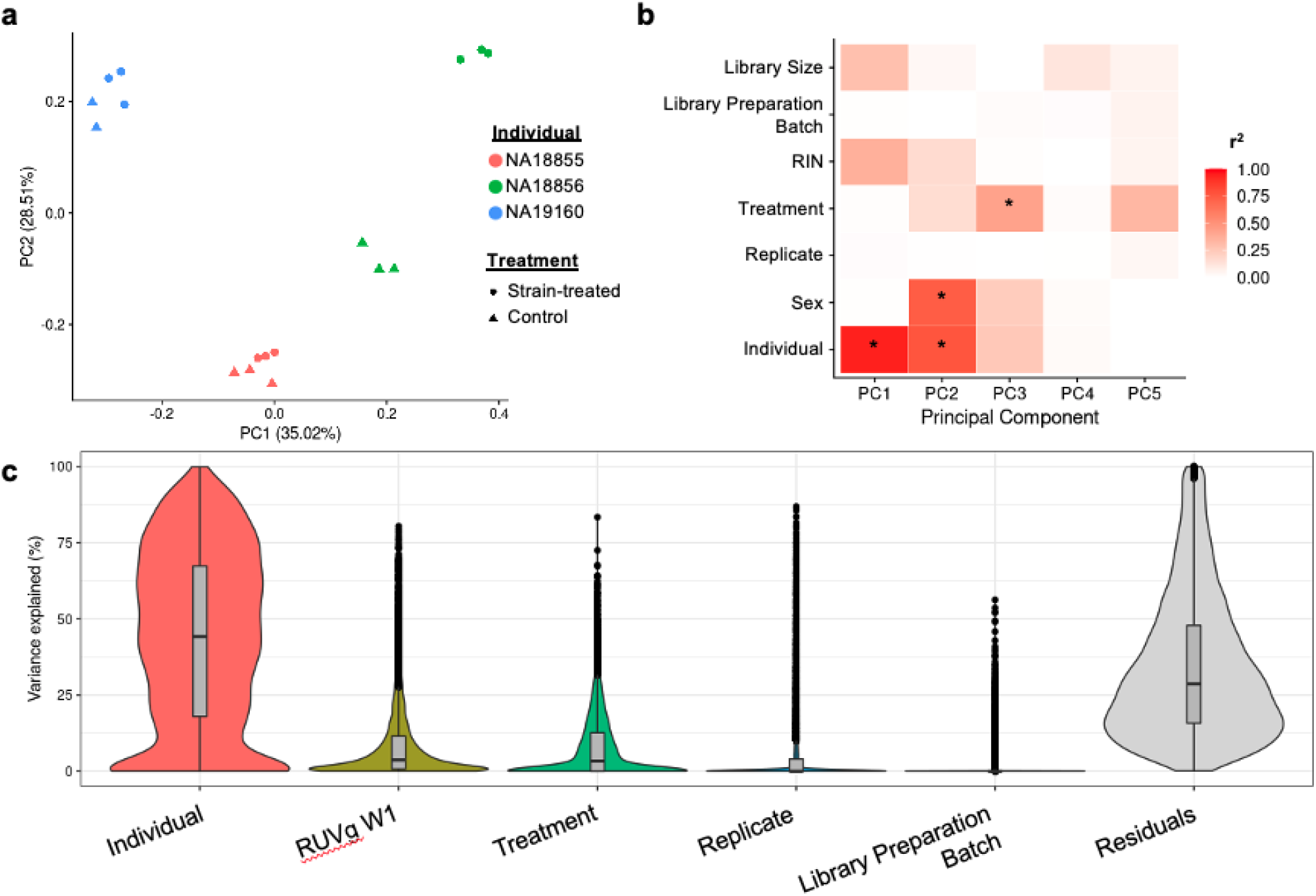
Characterization of sources of variation in bulk RNA sequencing data. **(a)** Principal components (PC) plot of normalized and RUVs-corrected bulk RNA sequencing samples colored by individual and shaped by treatment condition. Samples largely separate by Individual, with strain-treated samples from NA18856 showing a large separation from control samples from the same Individual. **(b)** Correlation between each of the first 5 PCs and several experimental variables, determined through linear regression analysis on normalized and RUVs-corrected bulk RNA sequencing data. Significant regressions (Benjamini-Hochberg corrected FDR < 0.05) are highlighted with an asterisk. **(c)** Violin-box plots displaying the fraction of variation explained by a number of experimental variables of the study design, including a single factor of unwanted variation fit using RUVg. Variables are ordered from largest to smallest by the median fraction of variation explained except for Residuals. The boxplots indicate the median, inner quartile range (IQR) and 1.5 times the IQR. Data beyond this are plotted as points. Violin plots indicate the density of data points based on their width.

Next, we took a more systematic approach to modeling the contribution of biological and technical factors to gene expression variation. Our goal was to leverage the total amount of variation in our data rather than focusing only on a few major axes of variation, as in the PCA above. We quantified the contributions of several experimental variables to gene expression variation on the level of individual genes using a linear mixed model (**Methods**). To do so, we modeled a single factor of unwanted variation in the data by using a set of 100 genes with the least amount of variation in the data as negative control genes^41^ (**Methods**). We then included the filtered and normalized gene expression data and this single factor of unwanted variation in the model (**Methods**; **Figure 5c**).

We determined that individual cell line contributes the largest amount of variance to the data (median of 42% variance explained). The additional factor of unwanted variation explains a median of 3.6% of the variance, and treatment explains a median of 3.5% of the variance. In contrast, technical replicate batch and cDNA library preparation batch explain a negligible amount of variance (median of 8.7×10^−7^ % and 3.5×10^−7^ % variance explained, respectively). We observed similar results when running a model that did not include the factor of unwanted variation (**Supplementary Figure S9**). Therefore, the biological variables of individual cell line and treatment contribute more to gene expression variation than technical variables. However, unwanted variation still seems to contribute to gene expression variation. Therefore, gene expression studies using this system should account for potential latent sources of variation.

### A power analysis

Our results are encouraging for our goal of verifying the feasibility of using iPSC-chondrocytes to study gene-by-environment interactions in OA. One possible way to study these interactions would be to use this system to map static eQTLs and dynamic response eQTLs (i.e., eQTLs that emerge in response to CTS). We conducted a power analysis to determine the potential impact of expanding this system to include 58 iPSC lines (**Figure 6**; we chose n=58 because this is the number of YRI iPSCs available to us). Under the assumptions of a simple linear regression to map eQTLs and a conservative Bonferroni correction for multiple testing (FWER = 0.05; **Methods**), we estimated that a sample of 58 individuals will provide 80% power to detect eQTLs with a standardized effect size of 0.7 in each of the control and treatment conditions. At this effect size, the power to detect eQTLs comes with a correspondingly low FDR (0.22).

**Figure 6:**
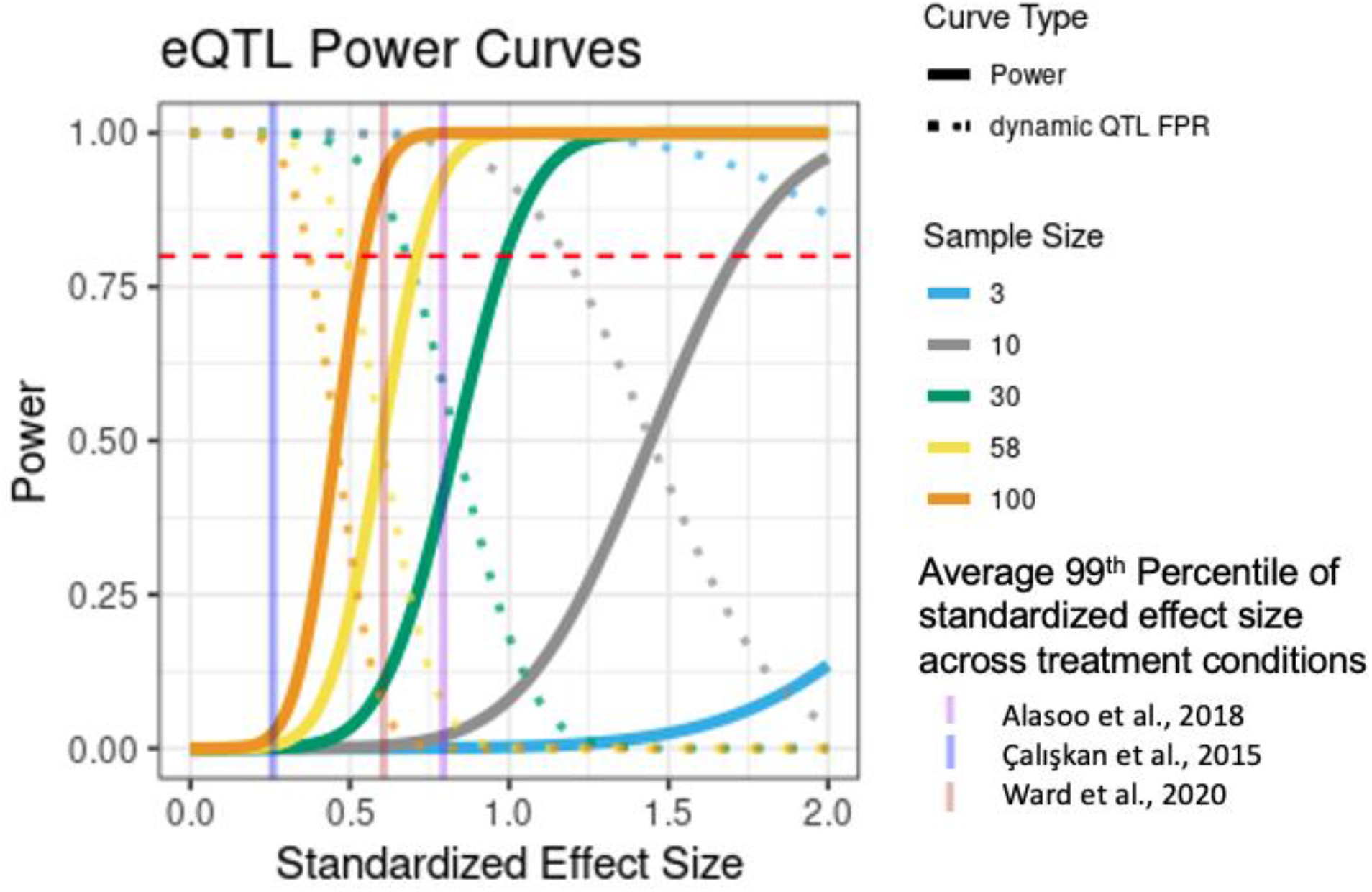
Power analysis for eQTL and dynamic eQTL study with two conditions. Power curves are derived under the assumption of a simple linear regression for eQTL mapping and plotted over standardized effect sizes (effect sizes divided by the phenotype standard deviation) for a range of sample sizes. Dynamic QTL false positive rates are computed as the probability of a SNP being called as significant in only one of two treatment conditions, assuming the standardized effect size was in fact identical in both conditions. The horizontal red line represents a power to detect eQTLs of 0.80. Vertical transparent lines represent the 99^th^ percentile of the standardized effect size estimated from an empirical cumulative distribution function fit to eQTL summary statistics from 3 published dynamic eQTL studies from different contexts, with the mean value over all the conditions in each study plotted.

To put these results into perspective, we reanalyzed eQTL summary statistics from a set of previous dynamic eQTL studies that come from a variety of different research contexts^12,42,43^ (**Figure 6**; **Table S7**). In each of these studies, hundreds to thousands of genes in each treatment condition have at least one eQTL which meets the standardized effect size threshold of 0.7 above. While none of these examples perfectly recapitulates the results of our *in vitro* OA system, the fact that these estimates are conservative and come from eQTL studies in three different stimulus conditions demonstrates the potential effectiveness of this approach.

## Discussion

We conducted a study to establish the feasibility of an *in vitro* iPSC-chondrocyte model for studying gene-by-environment interactions in OA. Gene-by-environment interactions, particularly those related to biomechanical stress, may play a role in OA pathogenesis. However, numerous ethical and logistical obstacles limit the study of these interactions and their effects on gene regulation in human chondrocytes. iPSC-chondrocytes may be a suitable alternative to circumvent these obstacles when paired with *in vitro* CTS models. Overall, our *in vitro* system allows for both the precise control of iPSC-chondrocyte environmental exposures and measurement of gene expression responses that are relevant to human joint health. While no *in vitro* model can completely accurately capture all aspects of *in vivo* disease, our results demonstrate that this system has tremendous potential to increase our understanding of OA and human joint health.

iPSC-chondrocytes are a valuable system to address the current lack of gene expression studies in human joint cells. Although iPSC-chondrocytes do not completely emulate mature, primary human chondrocytes, they do exhibit gene expression patterns characteristic of both adult chondrocytes and developing chondrocytes. The relatively early differentiation stage of our cells may be due to a variety of factors, including a shorter differentiation time and the culturing of cells as a monolayer as opposed to a 3-dimensional pellet^44^. Nevertheless, iPSC-chondrocytes provide a unique opportunity to learn about gene regulation in human joints and the basis of adult joint disease phenotypes. For instance, the ability to generate cells along the trajectory between iPSC-MSCs and mature chondrocytes allows for gene expression studies at a level otherwise infeasible with human primary tissues. Furthermore, prior studies have shown that studying iPSC-derived cells can uncover potentially important and transient forms of gene regulation that are masked in terminal cell types^10^.

Our results point to a robust gene expression response to CTS in iPSC-chondrocytes. We detected 987 DE genes in our study between treated and untreated cultures. These DE genes are enriched for gene sets relevant to joint health and OA. Thus, our results highlight the potential of this system as a platform for gene expression studies of human joint cells that enables research to circumvent the limitations of primary tissues. Our observations also suggest that studying gene regulation in this relatively simple system may provide insight into more complex biological processes relevant to human joint disease.

We acknowledge that *in vitro* CTS models do not directly mimic the compressive biomechanical stress felt by joint chondrocytes *in vivo*. However, CTS models are recognized as a valid method for studying the effects of extra-physiological stresses in cultured cells. Others have previously used CTS models to measure responses of joint cells to controlled biomechanical stress treatments^23–26,45–49^. Other groups have also developed models that use specific types and patterns of biomechanical strain to induce transcriptional and biochemical changes characteristic of early human OA^23–26^. Our results further attest to the utility of CTS as a model of biomechanical stress in studies of human joint health.

We also found that *in vitro* iPSC-chondrocyte system may be useful for studying the effects of genetic variation on gene regulation; moreover, it offers a way to study how biomechanical stress interacts with genetic factors to affect gene regulation. Indeed, individual-level differences drive a substantial amount of gene expression variation in this system. Therefore, eQTL and dynamic eQTL studies would be feasible using iPSC-chondrocytes. We identified specific differences in the gene expression response to CTS between individuals in this study. As such, iPSC-chondrocytes may be fruitful for uncovering gene-by-environment interactions involved in OA pathogenesis.

Investigating dynamic and context-specific gene regulatory effects may reveal the mechanisms contributing to OA development and progression, as this approach has been successfully applied to a variety of other cell type and trait contexts^9,12–16,42,50^. Previous studies have found that dynamic eQTLs are more enriched for relevant significant GWAS alleles than non-dynamic (‘standard’) eQTLs, which show consistent effects between conditions^9,10,13,15^. Our power analysis suggests that a study with a few dozen individuals may grant sufficient power to detect many static and dynamic eQTLs. Dynamic eQTLs may be more useful for identifying candidate susceptibility genes in joint diseases than steady state eQTLs, and they may also improve our understanding of gene-by-environment interactions related to OA and joint health.

Future studies using the iPSC-chondrocyte system should account for the possibility that transcriptional heterogeneity between and within individual iPSC-chondrocyte lines may confound association results in an eQTL study. Our analysis of the scRNA-seq data from control iPSC-chondrocytes suggests that differentiation efficiency does not differ substantially between individuals. Nonetheless, it is possible that differentiation efficiency may differ for other individuals not included in this study. There may also still exist transcriptional heterogeneity between iPSC-chondrocytes in their response to CTS that bulk RNA-seq would not adequately capture. Measuring and accounting for transcriptional heterogeneity in iPSC-chondrocytes twill also allow future gene expression studies to focus specifically on iPSC-chondrocytes, which represent only a minority of cells in each culture. This will increase power to detect both standard and dynamic eQTLs.

Our iPSC-chondrocyte system also opens the door to investigations beyond those involving only human cells. The existence of panels of human and nonhuman primate iPSCs^51^ introduces the possibility of inter-species comparisons of response to CTS. Comparative studies may help to uncover gene-by-environment interactions that contribute to the differential prevalence of OA and other joint diseases observed across primate species^52–54^.

We believe that the *in vitro* iPSC-chondrocyte CTS model shows great promise when applied to gene expression studies of OA. We hope such a system enables future studies of gene regulation in joint cells and their connections to joint health and disease.

## Methods

### Chondrogenic differentiation

iPSCs in this study were previously generated from transformed B cells derived from three Yoruba individuals (NA18855, female; NA18856, male; and NA19160; male) ^27^. Undifferentiated iPSCs were cultured on Matrigel-coated plates (Corning 356230) in Essential 8 (**E8**) medium at 37°C, 5% CO2, and atmospheric O2 until iPSCs reached 30% confluency. E8 medium was subsequently changed to mesenchymal stem cell (**MSC**) culture medium, which consisted of low glucose Dulbecco’s Modified Eagle Medium (**DMEM**) with 20% stem cell-qualified fetal bovine serum (**FBS**), and 100 mg/mL Penicillin/Streptomycin. The MSC medium was changed every day for 3 days, at which point cells were 80-100% confluent. On day 3, cells were detached from the Matrigel-coated petri dishes using a 0.05% Trypsin/EDTA solution and cultured on uncoated polystyrene flasks in MSC medium. The medium was changed every 2-3 days until the cells reached 90% confluency. The cells were then sub-cultured at a ratio of 1:3 until passage 4, at which point cells were classified as iPSC-derived MSCs. iPSC-derived MSCs were cryopreserved with cryopreservation media (80% FBS, 10% MSC culture medium, 10% Dimethyl Sulfoxide) in liquid nitrogen at passage 5 to 7.

iPSC-derived MSCs were detached from culture flasks using 0.05% Trypsin/EDTA and seeded at a density of 250,000 cells/well onto the center of wells of BioFlex Type I Collagen coated 6-well Culture Plates (FlexCell International BF-3001C) using BioFlex cell seeders (FlexCell International). Cells were seeded using a regimen of 15% elongation for 2 hours followed by overnight culture in MSC culture medium. After seeding, cells were cultured in serum-free chondrogenic differentiation medium^6^, consisting of high glucose DMEM, 100 mg/mL Penicillin/Streptomycin, 50mg/mL L-Proline, 200mM GlutaMax, 50mg/mL L-Ascorbic acid-2-Phosphate, 11g/L Sodium pyruvate, 5mM Dexamethasone, 1x ITS Premix, and supplemented with 10 ng/mL TGF-β3. The chondrogenic medium was changed every 2-3 days for 14 days.

### Standard phenotyping of iPSC-derived cells

Flow cytometry of iPSC-derived MSCs was performed using the BD Biosciences Human MSC Analysis Kit (BD Biosciences 562245), in combination with the Zombie Violet Fixable Viability Kit (BioLegend 423113). The Human MSC Analysis Kit assesses the surface markers CD90, CD105, CD73, CD34, CD45, CD11b or CD14, CD19, CD79α, and HLA-DR. In each flow experiment, matched iPSCs from the same line as each iPSC-derived MSC were included as a negative staining control. Samples were run on a BD LSRII Special Order System machine at the University of Chicago Cytometry and Antibody Technology Core Facility.

iPSC-chondrocytes were fixed using 4% paraformaldehyde in phosphate-buffered saline before staining using Alcian blue and Nuclear Fast Red. Alcian blue binds proteoglycans, which are found in connective tissue, particularly in cartilage^55^. Stained iPSC-chondrocytes and matched iPSC-MSCs from the same individuals were imaged using an Olympus dissecting microscope.

### Cyclic tensile strain regimen

iPSC-chondrocytes were treated with a cyclic tensile strain regimen that is known to induce an OA-like phenotype using the Flexercell FX6000 Tension System (Flexcell International)^23–26^. Plates were loaded onto the Flexercell baseplate (located in an incubator at 37°C, 5% CO2, and atmospheric O2), and a vacuum was used to deform the cell culture plate membrane and create uniform biaxial cyclic tensile strain. Specifically, 2.5% elongation (15kPa) of CTS was applied to the cells at a rate of 0.5 Hz for 24 hours.

### Droplet-based single cell RNA sequencing

iPSC-chondrocytes were dissociated from adherent conditions into single cell suspension as follows: First, cells were rinsed once with 1X PBS. Then, 1mg/mL of collagenase II) in 1X HBSS was added to cell culture wells at room temperature for 5 minutes. The collagenase II was neutralized with MSC medium and removed before further processing of the cells. Cells were rinsed once again with 1X PBS. A 0.25% Trypsin/EDTA solution was added to wells at room temperature for 2 minutes until cells detached. The trypsin was neutralized with MSC culture medium, and the cells were pelleted at 1000 rpm for 5 minutes and resuspended in FBS Stain Buffer. Cells were counted separately for each sample and combined in equal proportions before loading into a Chromium Single Cell A Chip kit (10X Genomics, 120236). To ensure that collection batch, individual, and treatment conditions were not confounded, samples were pooled strategically. One GEM well of a Chromium single cell chip targeting a collection of 5000 cells contained NA19160 control, NA18856 control, and NA18855 strain-treatment cells. A second GEM well of the same Chromium single cell chip targeting a collection of 5000 cells contained NA19160 strain-treatment and NA18855 control cells. The cells collected from sample NA18856 strain-treatment were not processed due to viability issues. Single cell cDNA libraries were established following the 10x Genomics Chromium Single Cell protocol^56^. In brief, the RNA of the captured cells was released by lysis, barcoded via a reverse transcription process, and amplified to produce gene expression libraries. The libraries were sequenced to 100 base pairs, paired-end on one lane using the Illumina HiSeq4000 at the University of Chicago Genomics Core Facility according to manufacturer instructions.

### Single cell data processing

FastQC was used to confirm that the reads were of high quality. Using an in-house computational pipeline, we extracted 10X cell barcodes and UMIs from raw scRNA-seq reads and mapped remaining reads to genes in the hg38 genome using STARsolo from the STAR software with default parameters (version 2.6.1b)^57^. The software *demuxlet* was used to deconvolute sample identity of individual cell droplets and detect multiplets in multiplexed samples with default parameters^58^. Previously collected and imputed genotype data for the three Yoruba individuals from the HapMap and 1000 Genomes Project were used as input for *demuxlet*^59,60^.

Processed gene count per cell barcode matrices were imported into R using the Seurat package (v3.2.0)^61,62^. Data were filtered to remove cells with fewer than 2000 UMIs detected and more than 10% of reads mapping to mitochondrial genes. Cells assigned as multiplets by *demuxlet* were also removed. A Uniform Manifold Approximation and Projection (**UMAP**) plot of the merged and unintegrated data shows that cells originating from the same individual cluster with each other (**Supplementary Figure S10**).

### Integration of individual level scRNA-seq data and characterization of cell clusters

Filtered scRNA-seq data was integrated across individuals using Seurat. Cells that were assigned as singlets by *demuxlet* were treated as individual datasets. Specifically, we focused in on just those datasets deriving from control cell culture conditions (n=3), as opposed to strain-treated conditions (n=2). Using Seurat, the *SCTransform* normalization function was applied to each of these datasets, and then datasets were integrated using integration anchors identified using the *FindIntegrationAnchors* function. Five-thousand features were selected as integration features for the SCT integration.

Seurat’s *FindClusters* function was used with 38 gene expression principal components (**PCs**) and a resolution of 0.4 as parameters to perform unsupervised clustering of transformed and integrated data. Thirty-eight gene expression PCs were chosen by locating the elbow in an elbow plot of PCs. To characterize the resulting three clusters that emerged, a Poisson adaptive shrinkage model was fit to the raw count data from the cells in each pseudo-sample described above using the ashR package ^63^. Poisson ashR models were fit separately for cell droplets assigned to each unsupervised cluster or separately for cell droplets from each individual. The cumulative density function of the inferred prior distributions for each of the fitted Poisson ashR models was plotted as in Sarkar and Stephens 2020^64^, for chondrogenic gene markers.

### Topic modeling of single cell RNA sequencing data

An unsupervised topic model with k=7 topics was fit to the scRNA-seq raw count data from several published sources and data from iPSCs and iPSC-derived cell types generated by our laboratory. Briefly, single cell data from iPSCs, iPSC-MSCs, iPSC-chondrocytes, and iPSC-osteoblasts collected by our group from a single human cell line were combined with single cell data from primary human hepatocytes^65^, iPSC-chondrocytes from an iPSC chondrogenic pellet time-course^66^, primary human chondrocytes^67,68^, and the iPSC-chondrocytes from the present study (details in **Supplementary Note**). Genes with non-zero counts in at least one cell in any of the six single cell datasets were included in the raw count matrix used to fit the topic model. A Poisson non-negative matrix factorization (**NMF**) model with 7 ranks was fit to the data using the *fit_poisson_nmf* function in the fastTopics R package with default parameters (v0.4.35)^69^. After fitting the Poisson NMF model, the fitted loadings and factors matrices were rescaled to sum to a total of 1 across each barcode for the loadings matrix and across each gene for the factors matrix to convert the Poisson NMF model into a topic model. The rescaled loadings matrix became the topic probabilities, and the rescaled factors matrix became the word probabilities in the resulting topic model.

The *diff_counts_analysis* function in fastTopics was applied to the topic model to evaluate differential expression of individual genes in each topic. Briefly, the function calculates a β statistic, which represents the log-fold change in relative occurrence of a gene in a single topic compared to its occurrence in all other topics. The function also calculates a standard error and z-score for each β statistic based on a Laplace approximation to the likelihood at the MLE.

### Bulk RNA extraction and sequencing

RNA was extracted from cells following CTS or control treatments using the ZR-Duet^™^ DNA/RNA MiniPrep kit (Zymo D7001). RNA concentration and quality were measured using the Agilent 2100 Bioanalyzer. Library preparation was performed over two batches using the Illumina TruSeq RNA Sample Preparation Kit v2 (RS-122-2001 & -2002, Illumina). Samples were sequenced to 50 base pairs, single-end on one lane using the Illumina HiSeq4000 at the University of Chicago Genomics Core Facility according to manufacturer instructions. A minimum of 17,284,094 raw reads were generated per sample. We used FastQC (http://www.bioinformatics.babraham.ac.uk/projects/fastqc/) to confirm that the reads were of high quality. One bulk RNA-seq sample was found to have a very low proportion of mapped reads (38.40%) and was excluded from subsequent analyses.

### Quantifying the number of bulk RNA-seq reads mapping to genes

Reads were mapped to the hg38 genome using STAR (version 2.6.1b)^57^. Gene expression levels were quantified using the *featureCounts* function in Subread (version 1.6.5) using standard parameters^70^. All downstream processing and analysis steps were performed in R (version 3.6.1) unless otherwise stated.

### Transformation and normalization of bulk RNA-seq reads

Log2-transformed counts per million (**CPM**) were calculated from raw counts for each sample using the edgeR package^71^. Lowly expressed genes were filtered such that only genes with an expression level of log2(CPM) > 2.5 in at least 4 samples were kept for downstream analyses. For the remaining 10,486 genes, the raw read counts were normalized using the relative log expression (**RLE**) method to account for the median number of reads sequenced across samples.

### Removing unwanted variation from bulk RNA-seq data

To account for batch effects arising between technical replicates before differential expression analysis, we modeled factors of unwanted variation using the RUVs correction method^41^ with k=2. RUVs is a method that uses technical replicate samples to estimate factors of unwanted variation from RNA-seq data. Individual-treatment pairs were constant within replicate blocks, which are used for the RUVs correction. RUVg is distinct from RUVs in that it uses negative control genes to estimate factors of unwanted variation from RNA-seq data rather than knowledge of technical replicate samples in the data.

### Differential expression analysis with bulk RNA-seq data

Differential expression (**DE**) was measured using a linear-model-based empirical Bayes method in the limma R package. The *voom* function from the limma package was also used to calculate weights to account for the mean-variance relationship in the RNA-seq count data.

Replicate batch was modeled as a random effect while treatment, individual, two RUVs coefficients, and RIN score were modeled as fixed effects in the linear mixed model for DE comparisons as in equation **(1)**. The ashR package^63^ was used to perform multiple testing correction on the DE tests using an adaptive shrinkage method. Genes with an FDR-adjusted p value < 0.05 were considered DE.

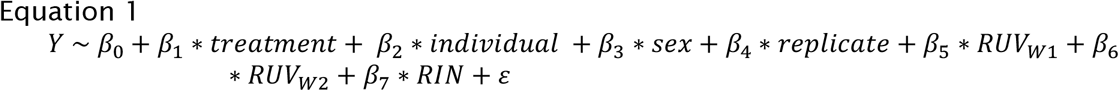

### Enrichment of DE genes in biological pathways and OA-related gene sets

Using topGO, we assessed enrichment of Gene Ontology (**GO**) biological processes among DE genes. A Kolmogorov-Smirnov test using ashR adjusted p-values was used for assessing enrichment of GO processes, and the top 20 most enriched terms were reported. To test for enrichment of sets of OA-related genes in our DE genes^2,4^, a Fisher’s exact test was used. In all enrichment tests, the background gene set was the complete set of genes tested for DE in our analyses.

There is the possibility that the enrichment of DE genes for GO categories and outside gene sets is driven by differential power due to some genes having higher expression in our samples. We therefore repeated the DE and enrichment analyses ten times, permuting the treatment condition labels for the samples each time.

### Analysis of sources of variation in bulk RNA-seq data

Principal component analysis (**PCA**) was performed on the normalized log2(CPM) values from above. A linear regression analysis was then performed between each of the top 5 PCs and several biological and technical variables. These variables included number of reads sequenced, library preparation batch, RNA integrity (**RIN**) score, treatment condition, replicate, and individual. P values from the regression were corrected using the Benjamini Hochberg (**BH**) procedure. Results with a BH-adjusted p value < 0.05 were considered significant.

The variancePartition package was applied to the filtered and RLE-normalized CPM values^72^. variancePartition uses a linear mixed model to quantify the contribution of variance from different sources. Our linear mixed model included variation due to individual cell line, treatment status, replicate batch, and library preparation batch. In addition, a single coefficient of unwanted variation was determined using the RUVg correction method^41^ with k=1; this coefficient was also included in the model. The RUVg correction method estimates factors of unwanted variation in RNA-seq data through negative control genes, which have the lowest variation in expression between samples. The 100 least variable genes in the data ranked by coefficient of variation were used as the set of control genes for the RUVg correction.

### Power curves for expression QTL (**eQTL**) and dynamic eQTL mapping

To ascertain the power to detect eQTLs and dynamic eQTLs across a range of sample sizes and standardized effect sizes, we followed the example presented in Ward *et al*., 2021^43^. In brief, for the power analysis we assumed a simple linear regression for eQTL mapping and a conservative Bonferroni correction for multiple testing (FWER=0.05). Standardized effect sizes are defined as the true additive effect size of genotype on gene expression divided by the phenotype standard deviation. To estimate the false positive rate of calling a dynamic eQTL, we computed the probability of a SNP being called as significant in only one of the two treatment conditions, assuming the standardized effect size was in fact identical in both conditions.

### Reanalysis of previous dynamic eQTL studies

We used summary statistics from eQTL mapping in three prior dynamic eQTL studies^12,42,43^ to determine standardized effect sizes for eQTL association tests in each treatment condition. Briefly, p values from association tests were converted into Z-scores using the appropriate quantile function. Z-scores were then converted to standardized effect sizes by adjusting for the square root of the sample size of the study. For summary statistics from Alasoo *et al*., 2018, and Caliskan *et al*., 2015, an adaptive shrinkage model was fit to the distribution of effect sizes and standard errors using ashR^63^. The ashR posterior estimates of effect sizes and standard deviations were used to compute the standardized effect size. Standardized effect size thresholds for at least 0.8 power under a sample size of 10, 30, 58, or 100 individuals were determined as described above. The number of genes with at least one association test that meets each of these thresholds in each condition were tabulated. Empirical distribution functions were fit to the distributions of the standardized effect sizes from each condition in each of the three studies. The 99^th^ percentile of these standardized effect sizes was determined from the empirical distribution function.

## Data availability

All computational scripts and analysis pipelines can be found on GitHub at https://github.com/anthonyhung/invitroOA_pilot_repository and in webpage format at https://anthonyhung.github.io/invitroOA_pilot_repository/index.html. All RNA-seq data have been deposited in the Gene Expression Omnibus (www.ncbi.nlm.nih.gov/geo/) under accession numbers GSE165874 and GSE167240.

## Acknowledgements

We thank Natalia Gonzales for comments on the manuscript. We thank Abhishek Sarkar, Michelle Ward, Kenneth Barr, and other members of the Gilad Lab for helpful discussions. We also thank Minal Caliskan for providing data files from her study “Host Genetic Variation Influences Gene Expression Response to Rhinovirus Infection.” This work was completed in part by using computational resources provided by the University of Chicago Research Computing Center.

## Author Contributions

G.H. conceived of the project with help from Y.G. and A.H. A.H., E.B., G.H., and C.G. performed the experiments. A.H. and G.H. performed the analyses. A.H. drafted the manuscript with input from Y.G. and G.H. Y.G. supervised the project.

