## SupplementaryInformation for "Characterizing gene expression responses to biomechanical strain in an *in vitro* model of osteoarthritis"

Supplementary Materials

**Supplementary Note** .......... 2
**Topic modeling external dataset details** .......... 2
**Supplementary Figures** .......... 3
**Figure S1: iPSC-MSC characterization.** .......... 3
**Figure S2: Alcian Blue staining of 14 day iPSC-chondrocytes and matched iPSC-MSCs.** .......... 4
**Figure S3: Additional gene markers of single cell clusters.** .......... 5
**Figure S4: Characterization of topics from topic model.** .......... 6
**Figure S5: PCA plots for normalized and filtered bulk RNA-seq data before RUVs correction including sample that failed QC.** .......... 7
**Figure S6: PCA plots for normalized and filtered bulk RNA sequencing data before RUVs correction.** .......... 8
**Figure S7: Correlation between each of the first 5 PCs and several experimental variables with normalized and non RUVs-corrected bulk RNA sequencing data.** .. 9
**Figure S8: PCA plots of additional PCs for normalized and filtered bulk RNA sequencing data after RUVs correction.** .......... 10
**Figure S9: Variance partition results on normalized and filtered bulk RNA sequencing data without the inclusion of RUVg factors of unwanted variation.** 11

### Supplementary Note

#### Topic modeling external dataset details

For the topic modeling analysis of scRNA-seq data, we included several external datasets in addition to the iPSC-chondrocytes from this current study.

First, scRNA-seq data collected by our group using the 10X Genomics Chromium Single Cell Gene Expression platform from iPSCs, iPSC-MSCs, iPSC-chondrocytes, and iPSC-osteoblasts all derived from a single human cell line were included in the topic modeling analysis. This human iPSC line was previously generated and characterized in our lab (Gallego Romero et al., 2015). The same protocol used to generate other iPSC-MSCs in this current study was also used to generate iPSC-MSCs here, with the exception that DMEM:F-12 (Thermo fisher 11330032) was used instead of low glucose DMEM (See **Methods)**. The same chondrogenic media formulation was used to differentiate iPSC-chondrocytes here as in this current study. iPSC-osteoblasts were generated by culturing iPSC-MSCs in osteogenic differentiation medium, consisting of high glucose DMEM (Gibco 11965092), 100 mg/mL Penicillin/Streptomycin (Corning 30002Cl), 10% stem cell-qualified fetal bovine serum (**FBS**, Thermo fisher 10567014), 50ug/mL Vitamin C, 100nM Dexamethasone, 10mM β-glycerophosphate, and 1uM Vitamin D. The osteogenic medium was changed every 2-3 days. Both iPSC-chondrocyte and iPSC-osteoblast protocols here included a total of 21 days of differentiation in their respective media before isolation and data collection, compared to 14 days for iPSC-chondrocytes in the current study.

Also included in the analysis were single-cell data from 3,490 hepatocytes published in MacParland *et al*., 2018 were subset from a larger dataset of single cell results from whole liver homogenate. Data from MacParland *et al*., 2018 are accessible using the R package HumanLiver and were originally obtained using the 10X Genomics Chromium Single Cell Gene Expression platform. These cells belong to clusters 1, 3, 5, 6, 14, and 15 identified in the original paper as showing enriched *ALB* (Albumin) expression, a hallmark of hepatocytes.

Additionally, data from iPSC-derived chondrocytes from a time-course of iPSC-chondrocyte pellet differentiation published in Wu *et al*., 2021 were obtained from GEO (GEO SRP290799). Data from single cells collected on day 7, day 14, day 28, and day 42 of differentiation were used to fit the topic model. Wu *et al*. chondrogenic pellets were treated with C59 for WNT inhibition during chondrogenesis to improve homogeneity of hiPSC chondrogenesis and avoid off-target cells.

Finally, data from 6,200 and 1,464 primary human chondrocytes were obtained from Chou *et al*., 2020 and Ji *et al*., 2018 respectively for use in the topic modeling analysis. Cells from Chou *et al*. 2020 were isolated from the intact outer lateral tibial plateau of a single male individual and processed using the 10X Genomics Chromium Single Cell Gene Expression platform. Data from these cells were downloaded from GEO (GEO Sample GSM4626766). Cells from Ji *et al*., 2018 were obtained from 10 patients with OA undergoing knee arthroplasty and underwent a modified single cell tagged reverse transcription (STRT) protocol for single cell transcriptional data collection. Data from all cells included in the original study were used. In both primary chondrocyte studies, isolated chondrocytes were not cultured *in vitro* before processing for scRNA-seq.

Although a model fit with seven topics (k=7) was ultimately chosen for further analysis, models fit with six or eight (k=6, k=8 respectively) topics did not differ substantially from this model (**Figure S11**).

### Supplementary Figures

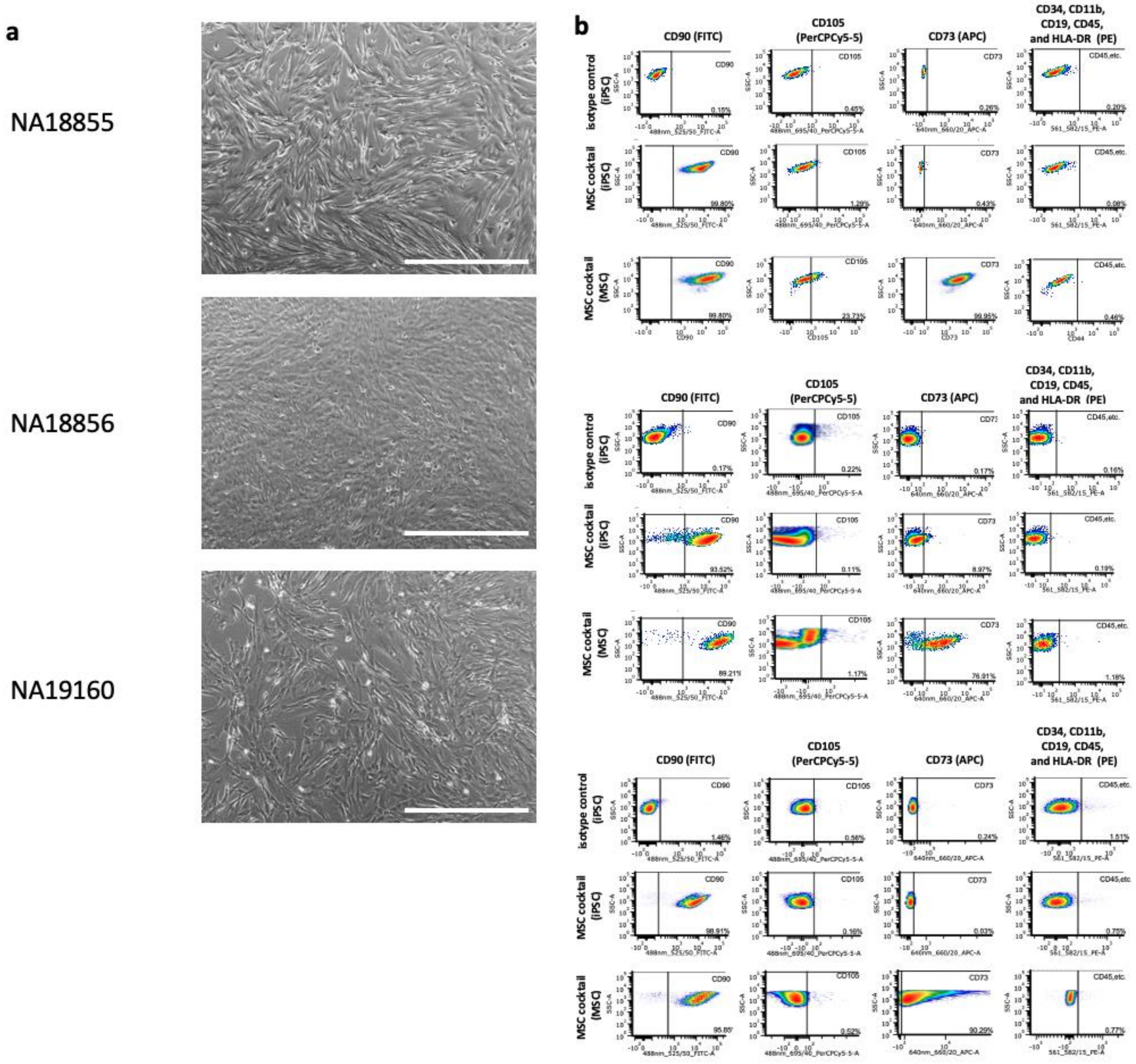

**Supplementary Figure S1: iPSC-MSC characterization.**
**(a)** EVOS microscope images of iPSC-derived MSCs from all three individuals demonstrate elongated morphology characteristic of MSCs. (Scale bar 250um) **(b)** Flow cytometry plots from labeling of iPSC-MSCs and matched iPSCs for cell-surface markers characteristic of MSCs. Compared to iPSCs, iPSC-MSCs demonstrate increased labeling of CD90 and CD73 while they do not show increased staining of several negative markers of MSCs. iPSC-MSC staining of CD105, a third positive marker of MSCs, is variable across the three individuals.

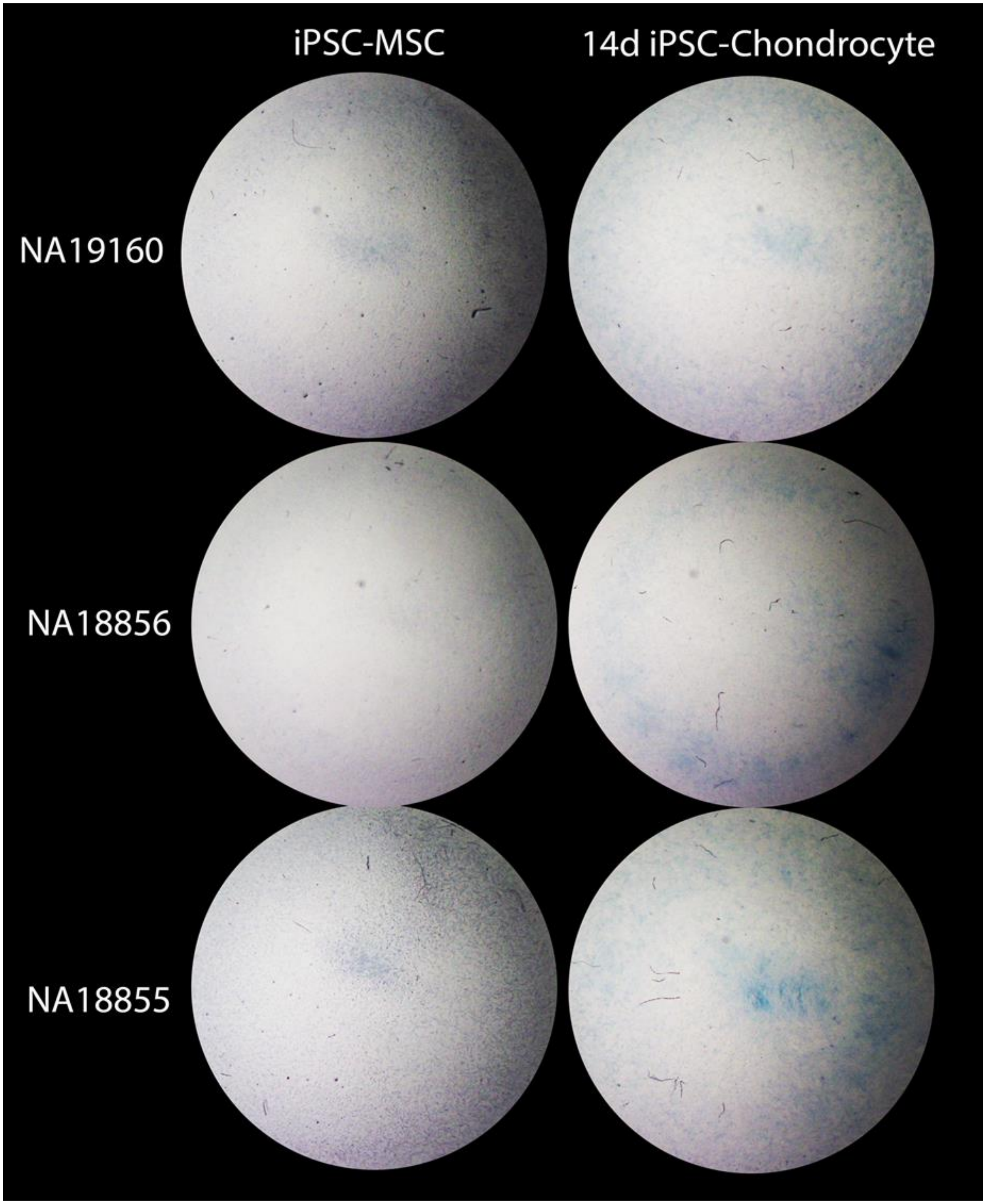

**Supplementary Figure S2: Alcian Blue staining of 14 day iPSC-chondrocytes and matched iPSC-MSCs.**

Brightfield dissecting microscope images are cropped to show circular central seeded area of cells on BioFlex Type I Collagen coated 6-well Culture Plates (diameter 25mm).

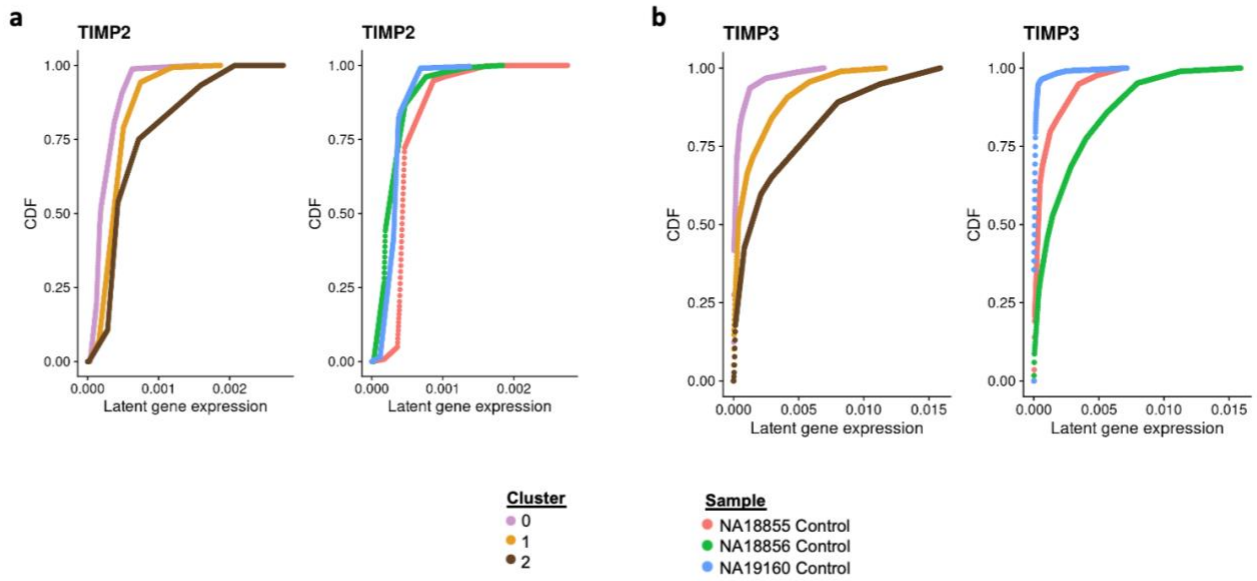

**Supplementary Figure S3: Additional gene markers of single cell clusters.** Cumulative distribution function (CDF) of marginal distributions of latent gene expression of **(a)** *TIMP2* and **(b)** *TIMP3* determined through fitting a Poisson adaptive shrinkage model to raw gene expression counts in each sample. In each panel, (Left) CDF curves colored by Seurat cluster. Cells in cluster 2 contain a higher latent gene expression of *TIMP2* and *TIMP3* on average. (Right) CDF curves colored by Individual.

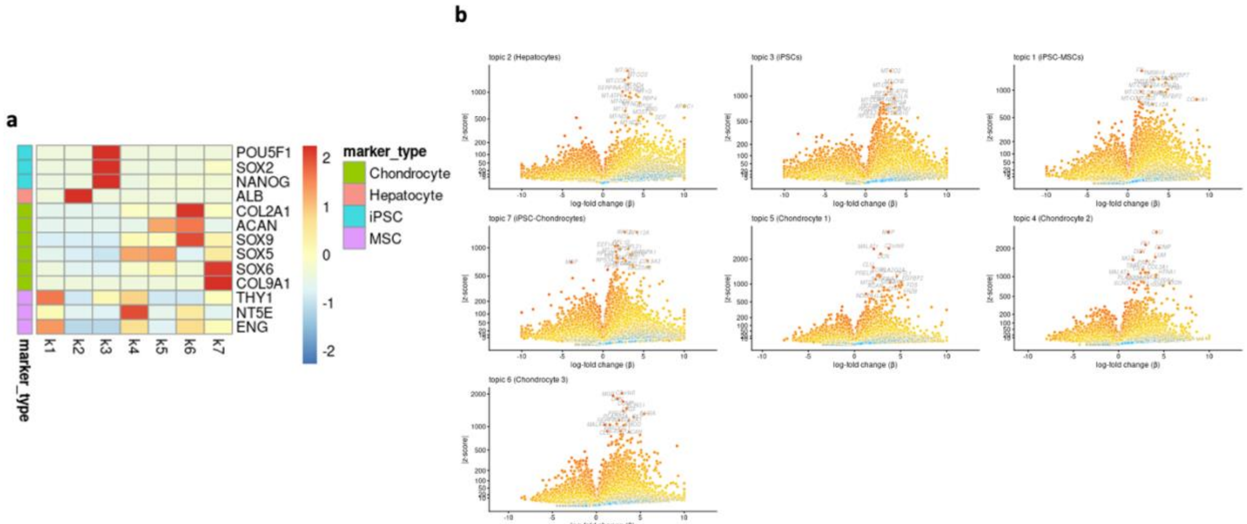

**Supplementary Figure S4: Characterization of topics from topic model.**
**(a)** Heatmap of word probabilities for selected marker genes for different cell types across a topic model containing seven topics. Heatmap cells are colored by word probabilities after applying a standard normal centering and scaling across each row. Colored bars on the left side of the heatmap denote the category of selected marker genes. Topic k2 demonstrates the highest word probability for hepatocyte marker gene *ALB*, while k3 demonstrates highest word probabilities for iPSC marker genes. Topic k1 demonstrates high word probabilities for MSC marker genes. Topics k4, k5, k6 and k7 demonstrate high word probabilities for several chondrocyte marker genes. **(b)** Volcano plots for log2 fold change relative occurrence (β) of individual genes in one topic compared to all other topics. |Z-scores| represent the absolute value of the z-scores for the β statistic. Positive β represent relative increase in occurrence of the gene in the topic compared to all other topics. Genes with |z-scores| above the 0.999 quantile are labeled in each plot.

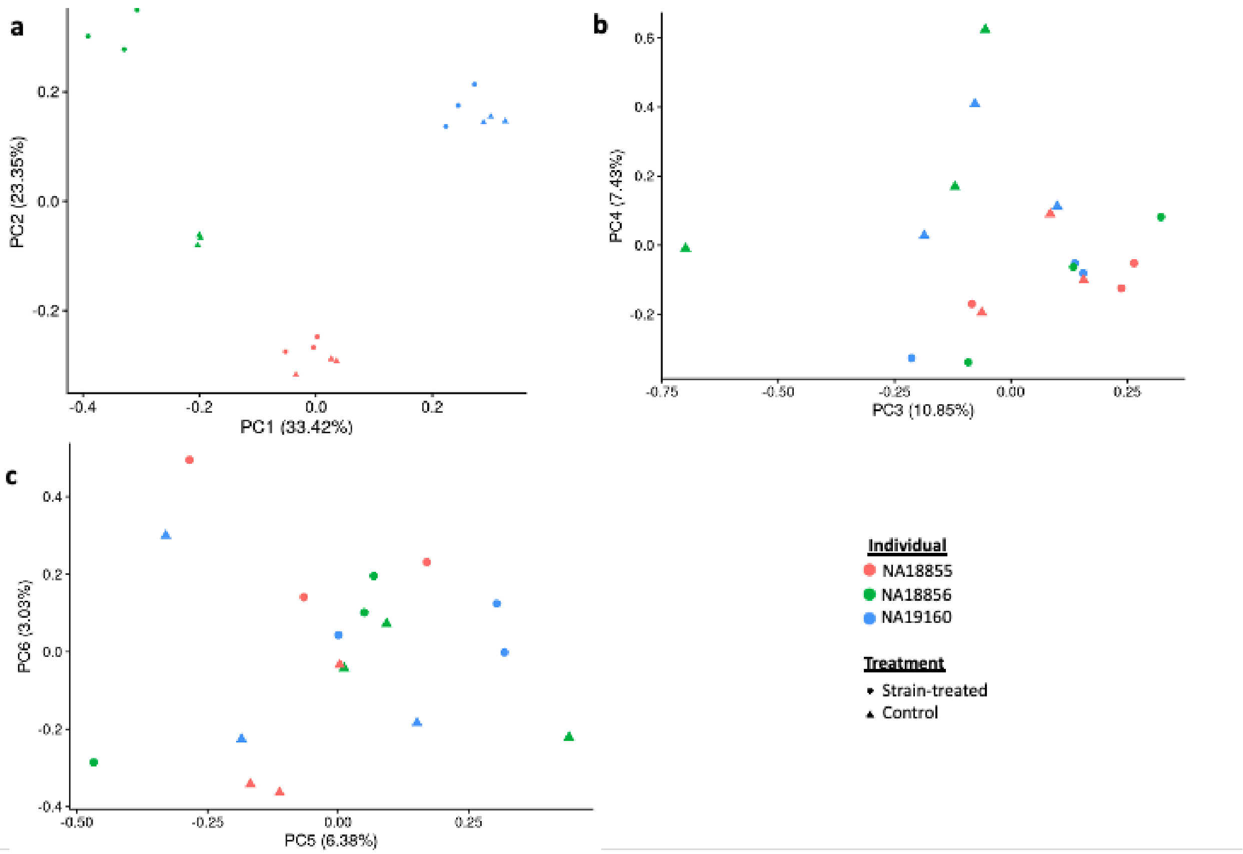

**Supplementary Figure S5: PCA plots for normalized and filtered bulk RNA-seq data before RUVs correction including sample that failed QC.**
**(a-c)** Principal components analysis (PCA) plots for principal components 1-6 are plotted for data including the single bulk RNA sequencing sample that was removed during quality control, corresponding to cells from individual NA19160 in the control condition from the first technical replicate of the experiment. When included in the PCA, this sample clusters as expected with corresponding samples from the other two technical replicates.

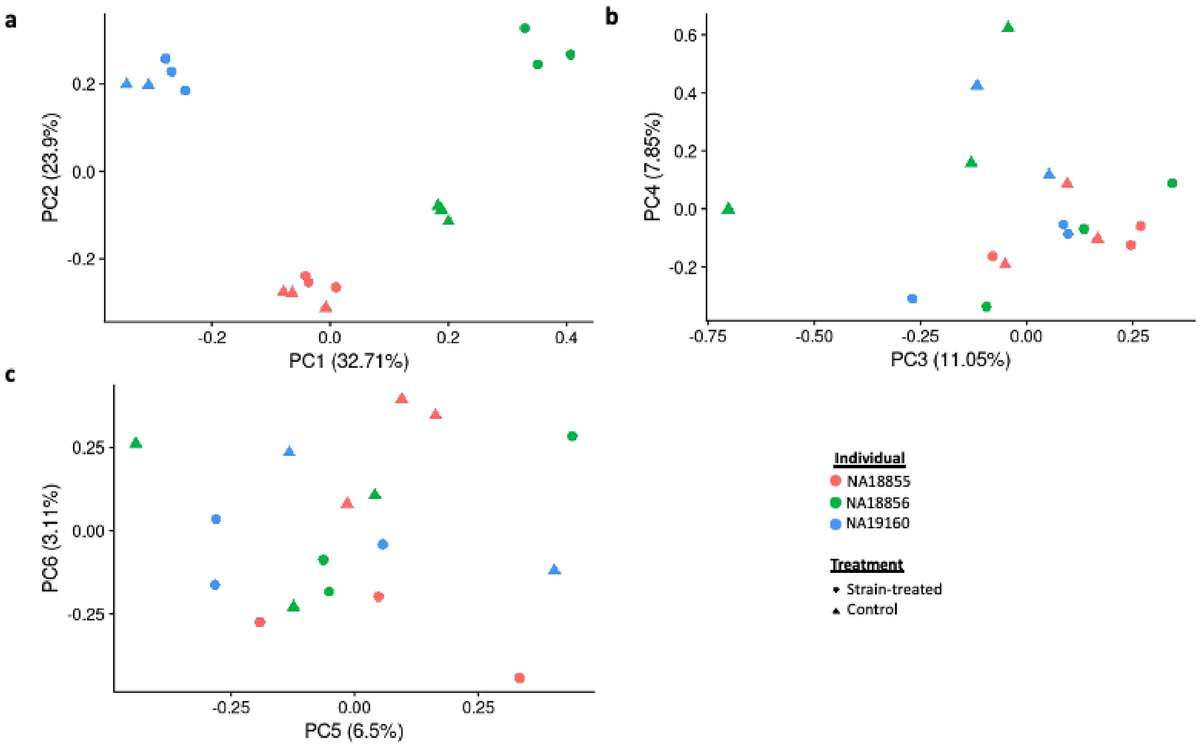

**Supplementary Figure S6: PCA plots for normalized and filtered bulk RNA sequencing data before RUVs correction.**
**(a-c)** PCA plots for principal components 1-6 are plotted for data absent RUVs correction.

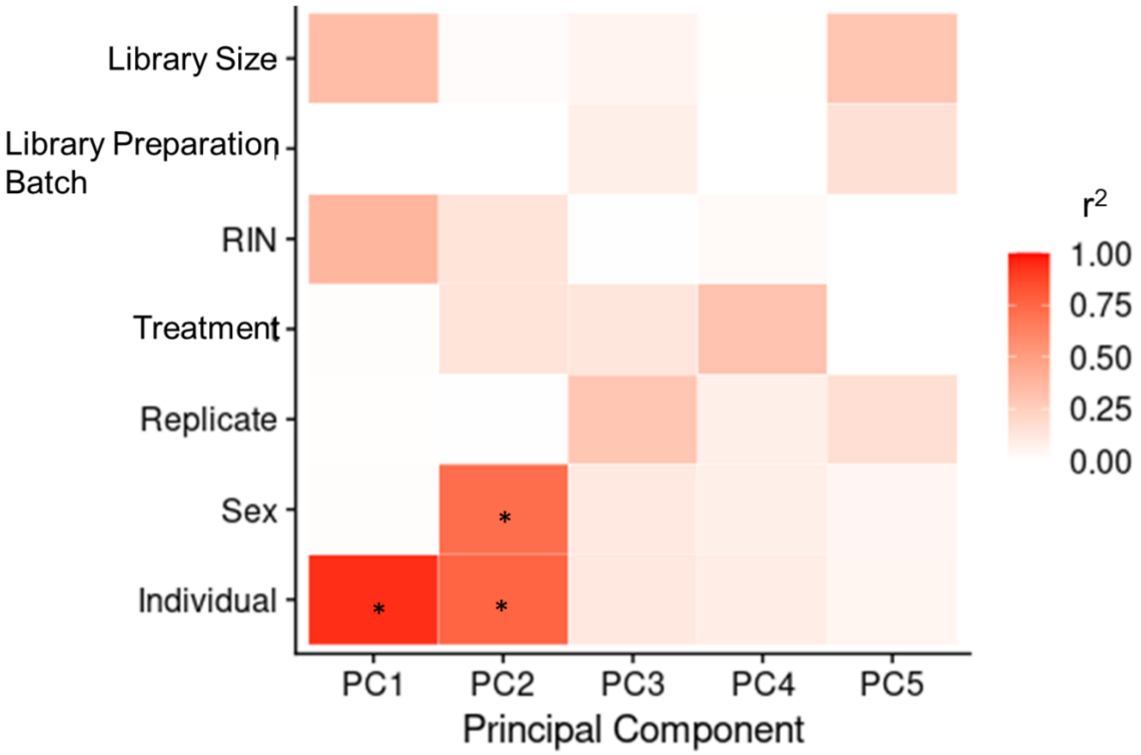

**Supplementary Figure S7: Correlation between each of the first 5 PCs and several experimental variables with normalized and non RUVs-corrected bulk RNA sequencing data.**
Significant regressions (Benjamini-Hochberg corrected FDR < 0.05) are highlighted with an asterisk.

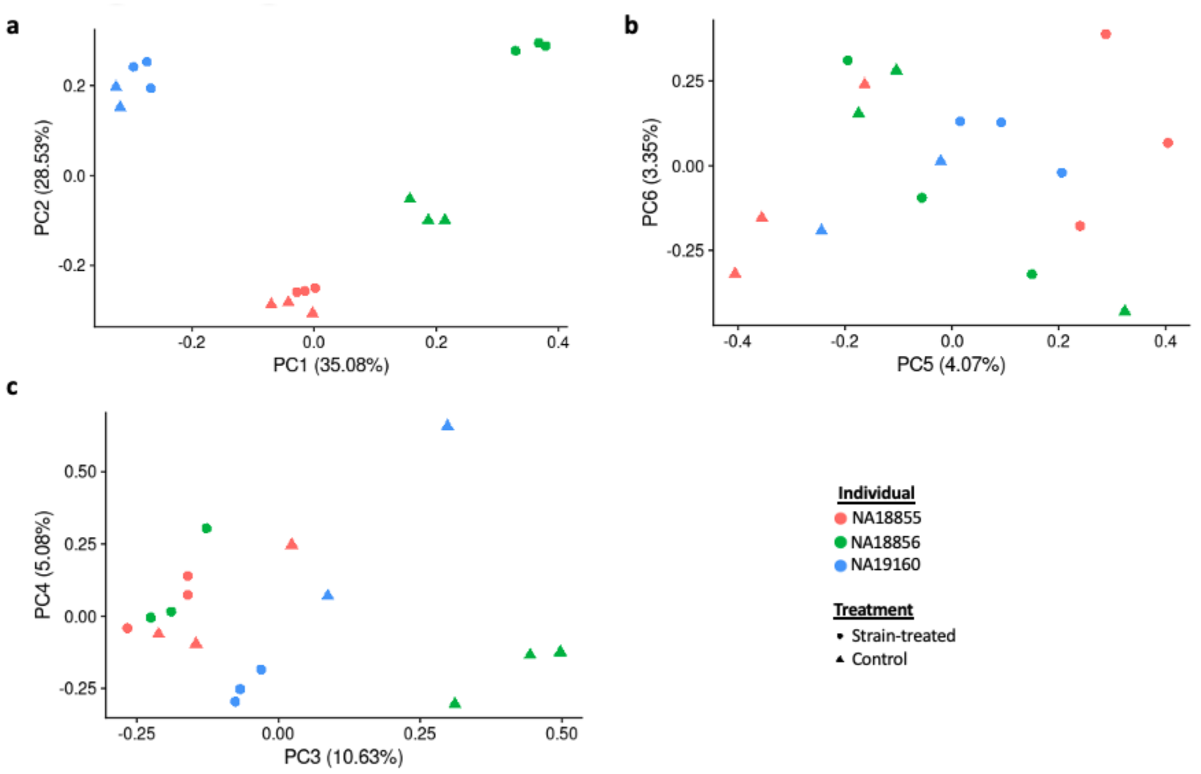

**Supplementary Figure S8: PCA plots of additional PCs for normalized and filtered bulk RNA sequencing data after RUVs correction.**

**(a-c)** PCA plots for PC3-6 for normalized and filtered bulk RNA sequencing data. PCA plot for PC1-2 is duplicated in Figure 3.

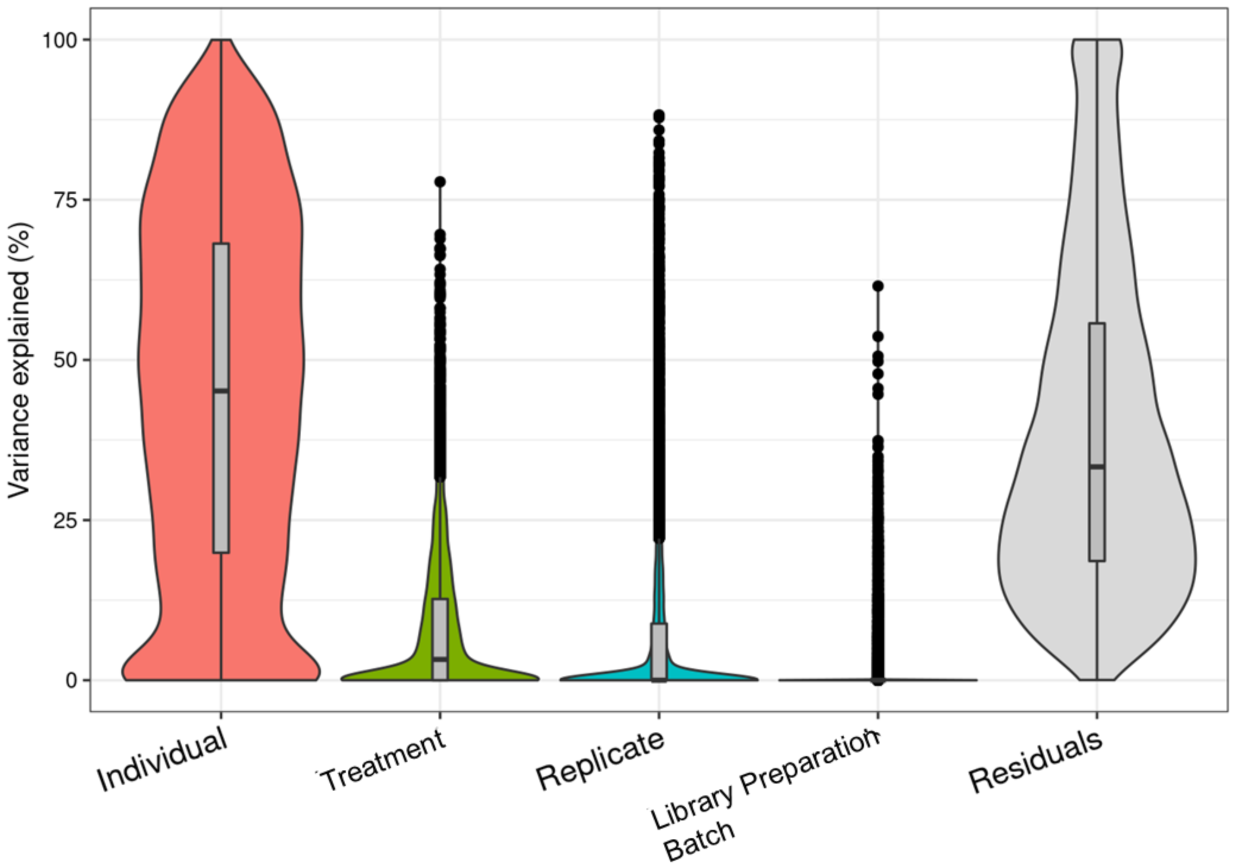

**Supplementary Figure S9: Variance partition results on normalized and filtered bulk RNA sequencing data without the inclusion of RUVg factors of unwanted variation.**
Individual and treatment both explain a larger proportion of variance of individual genes on average compared to technical factors of replicate and library preparation batch.

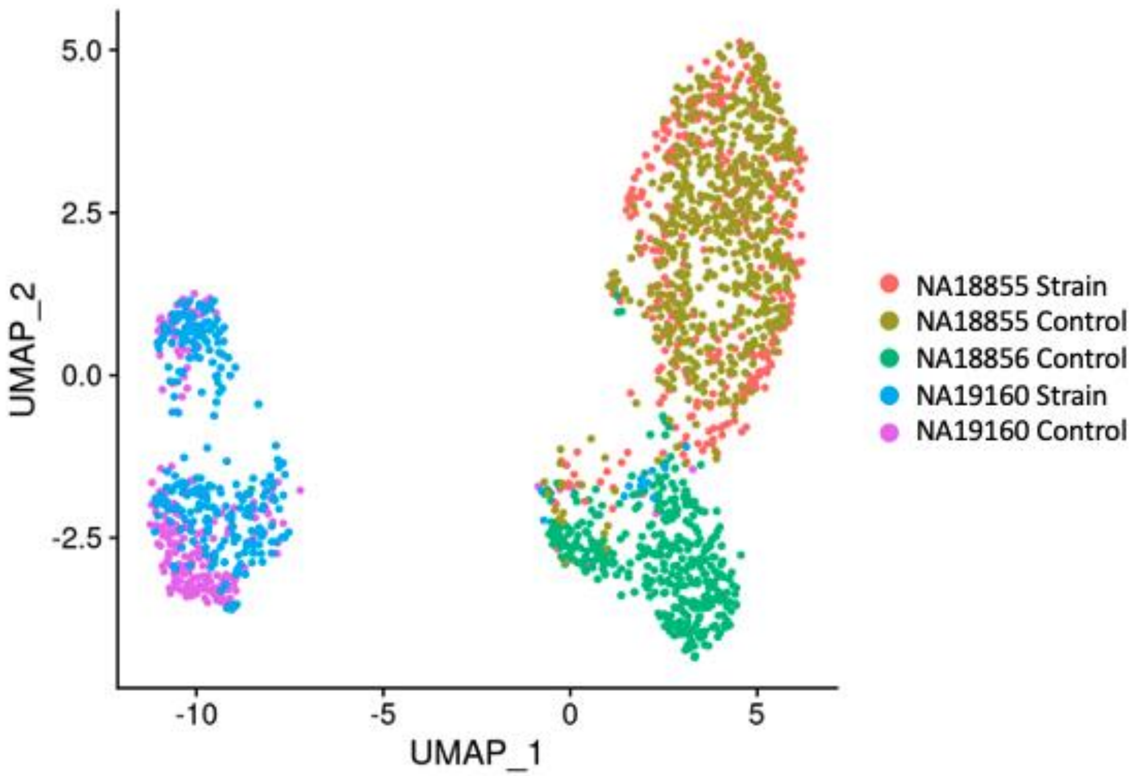

**Supplementary Figure S10: Uniform Manifold Approximation and Projection (UMAP) plot of merged scRNA-seq data.**
Raw scRNA-seq data from the two 10x GEM wells were merged and filtered to retain cells with fewer than 10% of reads coming from mitochondria and containing at least 2000 features. After log normalization, a UMAP plot was created for the data based on 2000 variable features and 50 gene expression principal components. No integration was applied to the data across the two GEM wells. Cells originating from the same individual cluster with each other.

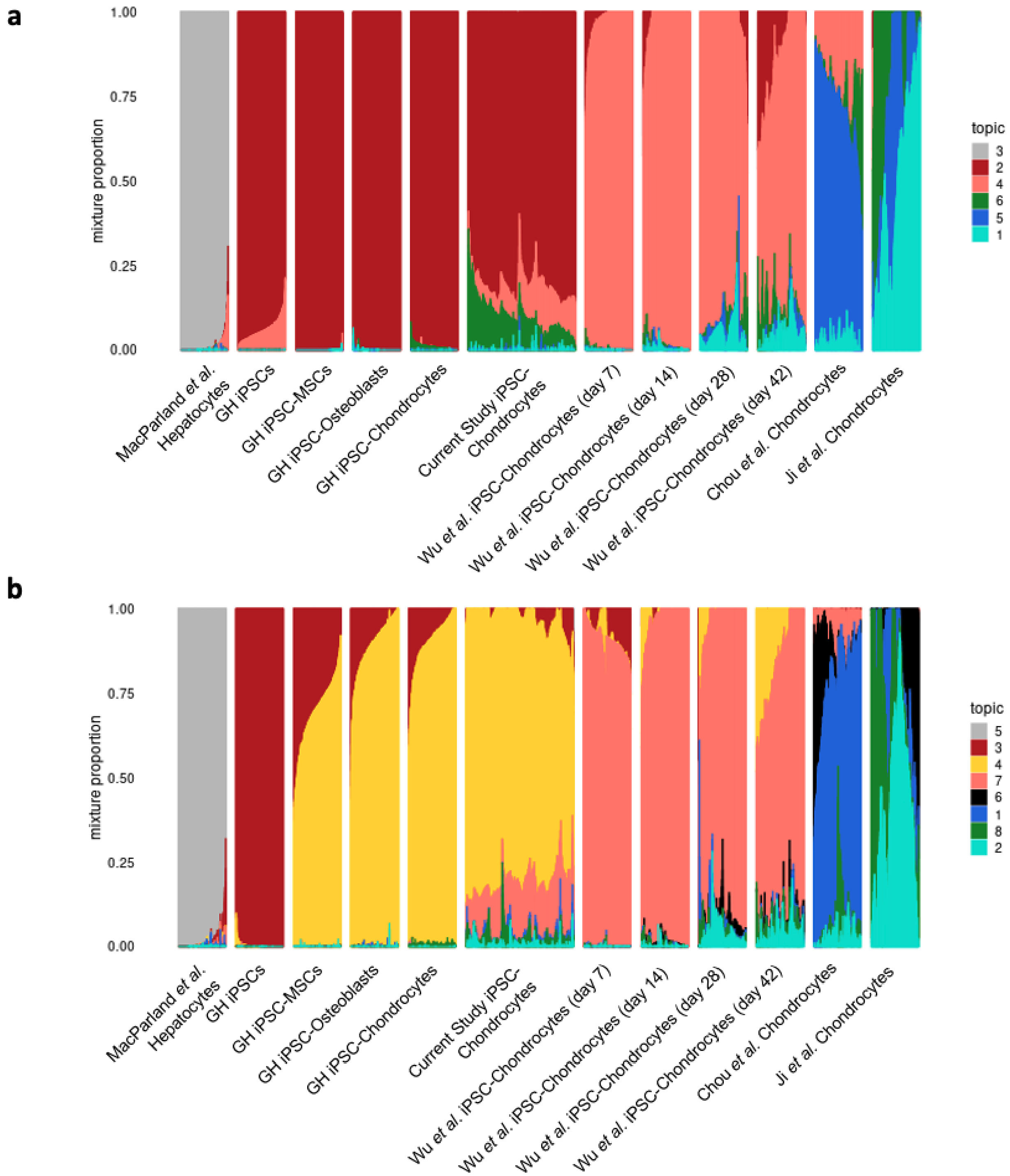

Supplementary **Figure S11: STRUCTURE plot for topic models fit to scRNA-seq data with k=6 and k=8.**
**(a)** k=6 **(b)** k=8 Colors representing topics are chosen to attempt to remain consistent between plots.

### Supplementary Table Legends

All supplementary tables are provided in Supplementary_tables.xlsx.

**Supplementary Table S1: Bulk RNA sequencing library metadata and quality control metrics.**

Sample_ID corresponds to a unique ID given to each of the 18 bulk RNA sequencing samples, corresponding to individual_replicate_treatmentCondition. Individual corresponds to the individual from which the iPSC-chondrocyte line was derived. Sex corresponds to the sex of the individual in the individual column. Replicate corresponds to the technical replicate of the experiment from which the sample is derived. Treatment corresponds to the treatment condition of the sample. RIN corresponds to the RNA integrity score of the bulk RNA library. Library Preparation Batch corresponds to the bulk RNA library preparation batch. Library Size corresponds to the library size of the bulk RNA library. cDNA Library Name corresponds to the unique ID label of the sample when submitted to the sequencing core. % Aligned corresponds to the percentage of reads that aligned to the human genome. M Aligned corresponds to the rounded million number of reads that were mapped. % Dups corresponds to the percentage of duplicated reads determined by STAR. % GC corresponds to the GC percentage of the reads in the sequenced library.

**Supplementary Table S2. Differential expression results from limma analysis between strain-treated and control samples.**

Each row corresponds to an independent test for a gene for differential expression, named by its Ensembl ID. logFC denotes the $\log_2$ fold change between control and strain-treated cells, with positive values indicating higher expression in control iPSC-chondrocytes. The AveExpr column gives the average $\log_2$-expression level for that gene across all samples. Column t denotes the moderated t-statistic for the test. Column P.Value lists the associated p-value for the test, and adj.P.Value is the p-value adjusted for multiple testing via a Benjamini-Hochberg procedure. ashr-lfsr, ashr-lfdr. ashr-qvalue, and ashr-svalue denote the ashR outputs for local false sign rate, local false detection rate, q value, and s value respectively.

**Supplementary Table S3. Genes from Tachmazidou *et al.*, 2019**

553 genes located within 500 kb of the 64 associated variants in – genes that were also identified by Tachmazidou *et al.*, 2019 as having prior evidence of involvement in animal models of skeletal disease or human bone diseases. These genes were determined by filtering rows from Supplementary Table 5 in the Tachmazidou *et al.*, 2019 paper by the "All_evidencee" column for any non-blank value.

**Supplementary Table S4. Genes with significant cross-omics differences between high-grade and low-grade cartilage in Steinberg *et al.* 2019.**

All external data in the table comes without modification from Supplementary Table 2 in the Steinberg *et al.*, 2019 paper.

**Supplementary Table S5. Results from 10 random permutations of sample treatment condition labels.**

Each row corresponds to a different permutation. The first column denotes the randomly assigned treatment condition label for each sample ID for a given permutation. The second column lists the top 20 enriched GO terms following DE analysis using limma/voom to fit a linear mixed model (equation 1) to the permuted data and assessing GO enrichment as described in the methods. The final two columns are the p-values for a Fisher's exact test for enrichment of permuted DE genes amongst the two sets of OA-related gene sets from Steinberg *et al.*, 2019 and Tachmazidou *et al.*, 2019.

**Supplementary Table S6. Full results of linear regression analysis between experimental variables and top 5 principal components in the bulk RNA sequencing data.**
Results from this analysis applied to normalized and filtered data before (top) and after (bottom) correction for 2 unwanted sources of variation using RUVs are reported. Columns in each table correspond to the first five principal components. Rows in each table correspond to several experimental variables. Values are either $r^2$ values for the correlations between the principal components and experimental variables as computed through a simple linear regression or Benjamini-Hochberg corrected q values for a test of association between the two variables.

**Supplementary Table S7. Results from reanalysis of prior dynamic eQTL studies for eQTL power analysis.**
Rows correspond to each of the treatment conditions from each of the three prior dynamic eQTL studies, colored by study. Columns correspond to the number of genes with at least one eQTL effect size in the treatment condition that reaches the effect size threshold for 80% power under an assumption of a sample size of 10, 30, 58, or 100.
